# PAST: latent feature extraction with a Prior-based self-Attention framework for Spatial Transcriptomics

**DOI:** 10.1101/2022.11.09.515447

**Authors:** Zhen Li, Xiaoyang Chen, Xuegong Zhang, Shengquan Chen, Rui Jiang

## Abstract

Rapid advances in spatial transcriptomics (ST) have revolutionized the interrogation of spatial heterogeneity and increased the demand for comprehensive methods to effectively characterize spatial domains. As a prerequisite for ST data analysis, spatial domain characterization is a crucial step for downstream analyses and biological implications. Here we propose PAST, a variational graph convolutional auto-encoder for ST, which effectively integrates prior information via a Bayesian neural network, captures spatial patterns via a self-attention mechanism, and enables scalable application via a ripple walk sampler strategy. Through comprehensive experiments on datasets generated by different technologies, we demonstrated that PAST could effectively characterize spatial domains and facilitate various downstream analyses, including ST visualization, spatial trajectory inference and pseudo-time analysis, by integrating spatial information and reference from various sources. Besides, we also show the advantages of PAST for accurate annotation of spatial domains in newly sequenced ST data and biological implications in the annotated domains.

---

Recent innovations in spatially resolved transcriptomics grant us a novel perspective on the cellular transcriptome, and hence have stimulated efforts to unravel cell transcriptomes in the context of cellular organizations and catalyze new discoveries in different areas of biological research^1-3^. When performing exploratory analysis of spatial transcriptomic (ST) data, one of the most critical steps is characterizing spatial domains^4^, which display spatially organized and functionally distinct anatomical structures on tissues. Nevertheless, characterizing spatial domains in ST data is computationally challenging, as the limitations of low capture efficiency and high dropouts in single-cell RNA sequencing (scRNA-seq) are inherited by most ST technologies^4^, resulting in sparse and highly noisy data. Besides, ST data usually represents substantial spatial correlation across tissues, requiring advanced approaches to take spatial localization information into account.

Several computational methods have been proposed to analyze ST data. BayesSpace performs spatial clustering via modeling the spatial correlation of gene profiles between neighboring spots^5^. SpaGCN identifies spatial domains by integrating gene expression, spatial location and histology through the construction of an undirected weighted graph^6^. CCST is an unsupervised cell clustering method based on graph neural networks for ST data^7^. STAGATE integrates spatial information and gene profiles with an adaptive graph attention auto-encoder to learn low-dimensional embeddings^8^. DR-SC utilizes a hidden Markov random field to perform dimension reduction and spatial clustering within a unified framework^9^. Besides, Scanpy and Seurat, two widely-used single-cell analysis workflows in Python and R communities, respectively, also provide vignettes tailored for ST data analysis^10, 11^.

However, effective approaches for characterizing spatial domains in ST data need to overcome several challenges. First, existing methods failed to incorporate prior information from reference data to facilitate ST data analysis. Incorporating prior information in analyzing single-cell genomic data can better tackle the high level of noise and technical variation, and has been successfully applied in the analysis of various single-cell genomic data^12-15^. With the advanced technologies, massive amounts of ST data have been generated by consortiums or individual research groups^16-18^ and accumulated in repositories^3, 4, 19, 20^, paving the way to taking full advantage of the existing data and leveraging the data as reference to characterize spatial domains. Second, local spatial patterns should be effectively captured. Although it has been observed that aggregating information from each spot’s neighbors via a graph convolutional network (GCN) is a promising approach to identify localized gene expression patterns^4, 6, 7^, tailored model design for better characterization of local spatial patterns is still demanded. Third, global spatial patterns should be carefully preserved. With the exponential growth of profiled spots and genes, the graph-based approach becomes cumbersome and time-consuming, requiring strategies that perform memory and time-efficient mini-batch training and prediction while preserving global spatial patterns. Fourth, there is an emerging need for the automatic annotation of spatial domains, since advanced protocols constantly increase sequencing throughput, and the typical approach, which performs unsupervised spatial clustering and then assigns the putative domain to each cluster subjectively, would be cumbersome and irreproducible for large-scale ST data.

To this end, we propose PAST, a **P**rior-based self-**A**ttention framework for **S**patial **T**ranscriptomics, to effectively characterize spatial domains. PAST is built upon a variational graph convolutional auto-encoder, and utilizes a Bayesian neural network (BNN) to incorporate prior information from reference data, self-attention mechanism^21^ to capture local spatial patterns, and ripple walk sampler^22^ to enable scalable subgraph-based training and prediction. The latent embeddings obtained from PAST contain valuable spatial correlation patterns and can be directly fed into existing scRNA-seq computational tools for effective and novel downstream analyses. Through comprehensive experiments on multiple datasets from different ST technologies, we demonstrated that PAST outperforms baseline methods for spatial domain characterization, ST visualization, spatial trajectory inference, and pseudo-time analysis. Besides, PAST could effectively leverage reference data from various sources and provide satisfactory robustness and scalability. In addition, we showed that PAST not only opens a new avenue for automatically annotating spatial domains of newly sequenced ST data, but also has the potential to provide biological insights into the annotated domains.

## Results

### Overview of PAST

PAST is a weakly supervised model based on a framework of variational graph convolutional auto-encoder, taking transcriptomic profiles and spatial coordinates of the target ST data and transcriptomic profiles of the reference data as input. As shown in Fig. 1, the first layer of the encoder decomposes the total variation in ST data into two components: the component that captures the shared biological variation with reference data and the component that captures the unique biological variation in target ST data. More specifically, the first component utilizes a BNN to incorporate prior information of the projection vectors learned from reference data, and it captures the variation in target ST data that is shared with the reference data. Choice of the reference data is flexible and can be constructed from various sources: (1) other ST data of the same tissue and from the same protocol with the target ST data, (2) other ST data of similar tissues and from other protocols, (3) scRNA-seq data of similar tissues, or (4) the target ST data itself. In practice, the reference data can be incomplete: novel domains of biological variation that are not captured in the reference data can be present in the target ST data. The second component utilizes a fully-connected neural network to adaptively capture the unique variation in target ST data that is not present in reference data (Methods).

**Fig. 1.**
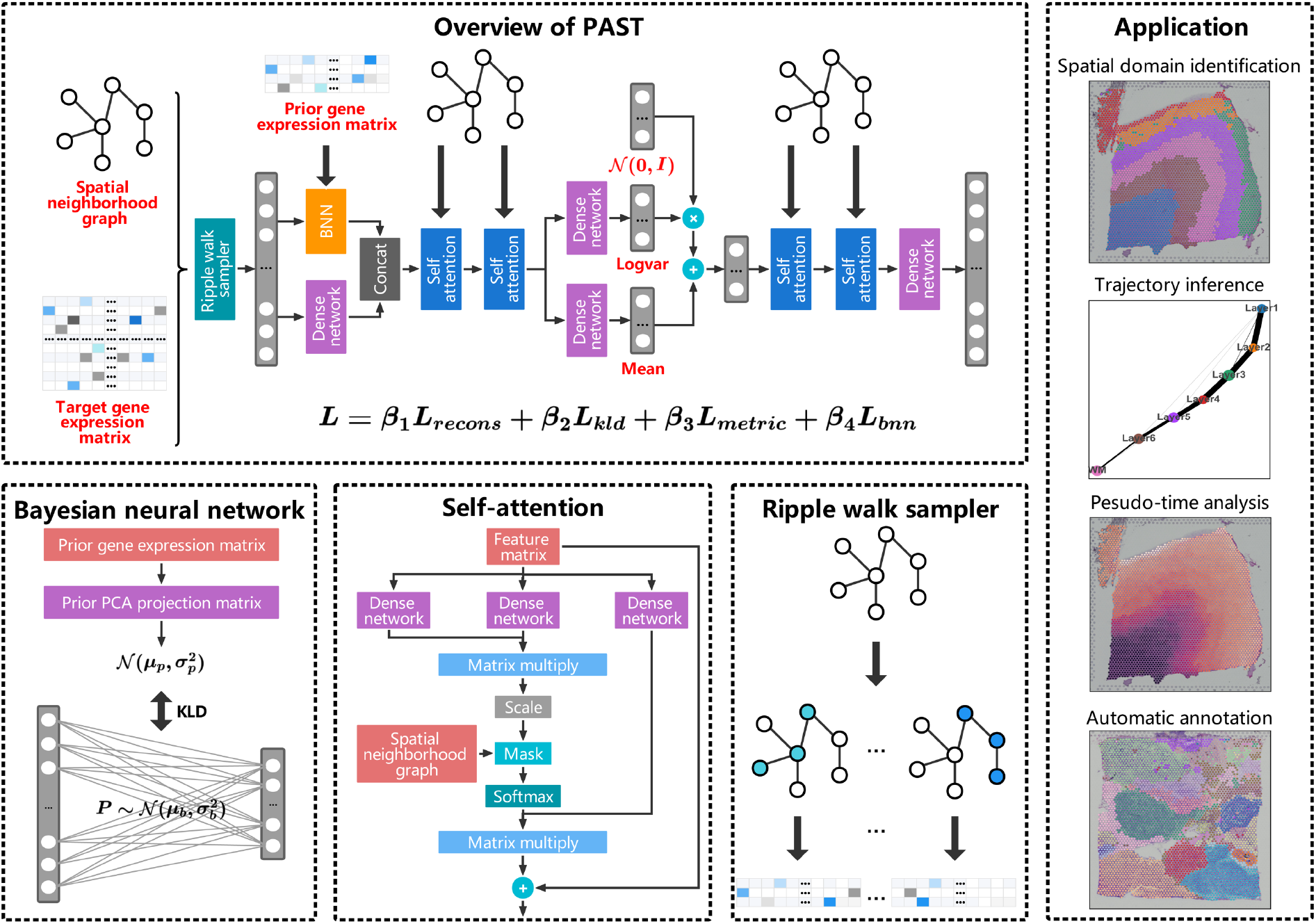
The PAST framework. PAST is built on a variational graph convolutional auto-encoder, taking transcriptomic profiles and spatial neighborhood graph of target data and transcriptomic profiles of reference data as input. PAST integrates shared biological variation of reference data from various sources based on BNN, effectively captures spatial correlation between neighboring spots via the self-attention mechanism in graph convolutional networks, and samples high-quality subgraph satisfying connectivity and randomness for scalable training and prediction based on a ripple walk sampler. After model training, PAST outputs the embeddings of cell in the target ST dataset and facilitates various downstream tasks including spatial domain identification, trajectory inference, pseudo-time analysis, and automatic domain annotation.

To integrate information of transcriptomic profiles and spatial coordinates, PAST identifies *k* nearest neighbors (*k*-NN) for each spot using spatial coordinates in a Euclidean space, and adopts GCNs to aggregate spatial patterns from each spot’s neighbors. Specifically, the self-attention mechanism^21^, which fits the relationship between words well in machine translation tasks^23^, is used to model local spatial patterns between neighboring spots, while the ripple walk sampler^22^, which enables efficient subgraph-based training for large and deep GCNs^24^, is used to achieve better scalability on large-scale ST data and preserve global spatial patterns simultaneously. PAST also restricts the distance of latent embeddings between neighbors through metric learning^25^, the insight of which is that spatially close spots are more likely to be positive pairs to show similar latent patterns (Methods). After model training and prediction, the latent embeddings can be applied to spatial domain characterization, spatial visualization, trajectory inference, pseudo-time analysis, and automatic domain annotation.

### PAST effectively incorporates reference data and characterizes spatial domains

We first use an example as a proof of concept to demonstrate PAST. We collected 12 human dorsolateral prefrontal cortex (DLPFC) sections from a 10x Genomics Visium dataset^26^. For each section, we constructed the reference data from the ST data itself and from external ST data, namely the other 11 sections (Methods). The two variants of PAST have different application scenarios and are referred to as PAST-S and PAST-E, respectively. PAST was benchmarked against seven baseline methods, including two widely-used single-cell analysis workflows (Scanpy^10^ and Seurat^11^), and five methods for ST data analysis (SpaGCN^6^, BayesSpace^5^, CCST^7^, DR-SC^9^, and STAGATE^8^). Note that the vignettes tailored for ST data analysis and the default parameters or settings of the baseline methods were used (Methods). We demonstrate the advantages of PAST from the following four perspectives.

First, we quantitatively evaluated the performance of spatial domain characterization via (1) supervised cross-validation, (2) unsupervised spatial clustering with a specified number of clusters, and (3) unsupervised spatial clustering with default resolution. The supervised cross-validation aims to assess the representation capabilities of the latent embeddings learned by various methods. Specifically, we treated the latent embeddings as the input and the labels of spatial domains as the output, adopted support vector machine (SVM) as the classifier as suggested by benchmark studies on supervised single-cell annotation^27, 28^, and conducted 5-fold cross-validation experiments (Methods). We evaluated the performance by the average score of accuracy (Acc), Cohen’s kappa value (κ), mean F1 score (mF1), and weighted F1 score (wF1), respectively, in the 5-fold experiments (Methods). A higher score of the metrics suggests that the latent embeddings can better predict the spatial domains and, consequently, have superior representation capabilities. As shown in Fig. 2a and Supplementary Fig. S1a, PAST-S and PAST-E significantly outperformed the baseline methods. One-sided paired Wilcoxon signed-rank tests showed that PAST-S and PAST-E achieved better performance than STAGATE, the most recent spatial embedding method, with *p*-values of 0.0046 and 0.0017 for accuracy, respectively (0.0024 and 0.0012 for κ, 0.0005 and 0.0005 for mF1, and 0.0007 and 0.0005 for wF1), indicating the effective spatial domain characterization.

**Fig. 2.**
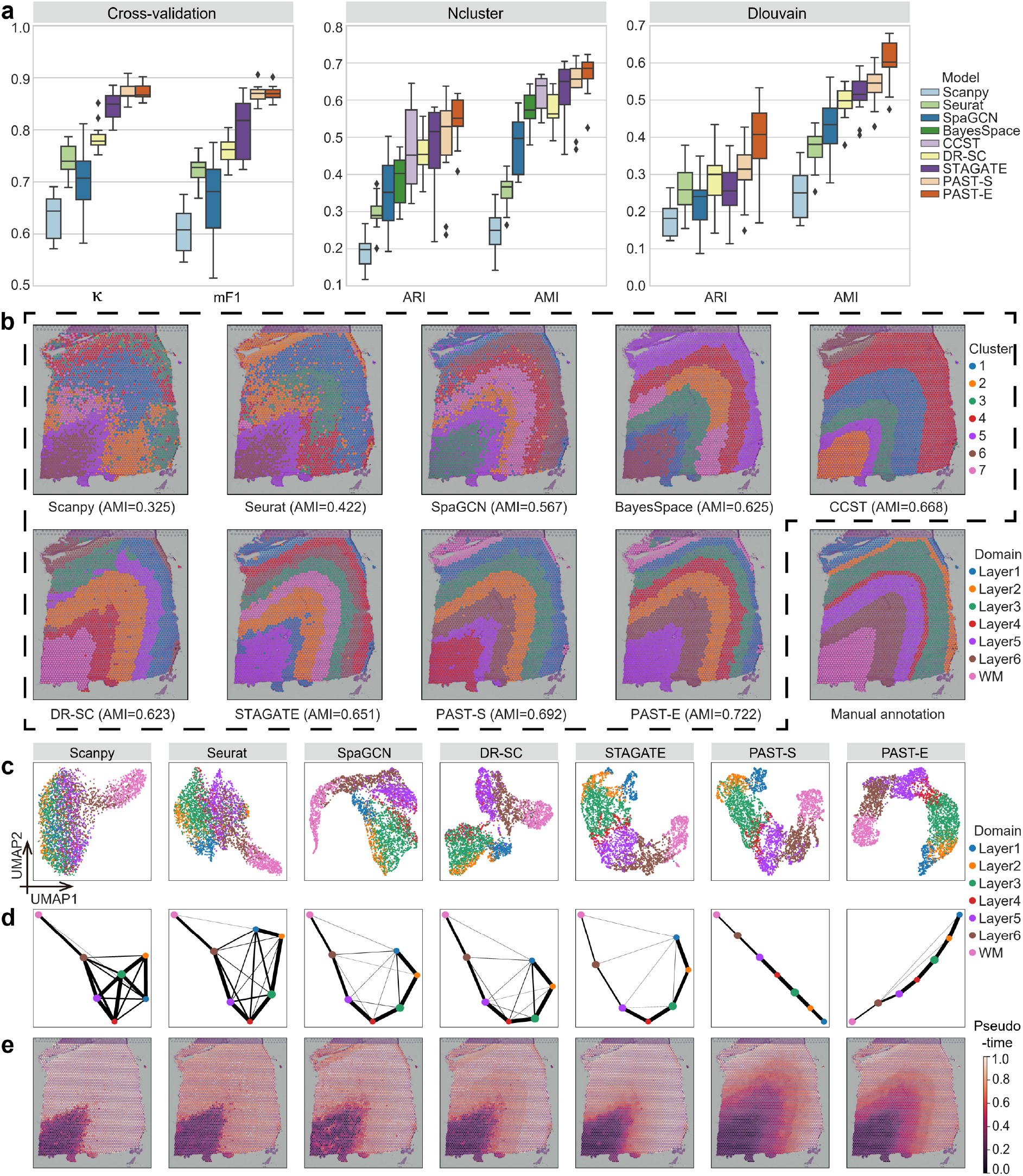
PAST effectively incorporates reference data and characterizes spatial domains. **a**, Quantitative performance evaluation of spatial domain characterization on 12 DLPFC slices via supervised cross-validation and unsupervised spatial clustering with a specified number of clusters (Ncluster) and with default resolution (Dlouvain). The cross-validation performance was evaluated by the average score of Cohen’s kappa value (κ) and mean F1 score (mF1), respectively, in the 5-fold experiments. The spatial clustering performance was evaluated by adjusted rand index (ARI) and adjusted mutual information (AMI). The center line, box limits and whiskers in the boxplots are the median, upper and lower quartiles, and 1.5× interquartile range, respectively. **b**, Visualization of the results of spatial clustering with a specified number of clusters and the manual annotation of Slice 151673. **c**, UMAP visualization, **d**, PAGA trajectory inference results and **e**, DPT pseudo-time analysis results of Slice 151673.

For the unsupervised spatial clustering with a specified number of clusters, we directly utilized the baseline spatial clustering methods, such as BayesSpace^5^ and CCST^7^, or the accompanying spatial clustering strategy of each of the other baseline methods on the learned latent embeddings. Following STAGATE, PAST performed spatial clustering with a specified number of clusters by the mclust algorithm. For each DLPFC section, the number of clusters was specified as the number of unique ground-truth spatial domains. For Scanpy and Seurat which perform clustering by Louvain or Leiden algorithm, we implemented a binary search to tune the resolution parameter in clustering to make the number of clusters and the specified number as close as possible (Methods). We evaluated the clustering performance by six metrics: adjusted rand index (ARI), adjusted mutual information (AMI), normalized mutual information (NMI), fowlkes-mallows index (FMI), completeness (Comp), and homogeneity (Homo) (Methods). PAST-S and PAST-E again provided superior performance than the baseline methods (Fig. 2a and Supplementary Fig. S1a). Scanpy and Seurat, two widely-used single-cell analysis workflows, provided relatively poor performance even using the vignettes tailored for ST data analysis, indicating the importance of taking full advantage of the spatial information. Note that for BayesSpace and CCST, we only evaluated the clustering performance with a specified number of clusters because they are spatial clustering methods and do not provide latent embeddings.

For unsupervised spatial clustering with default resolution, which is much more useful for the general situation where the number of unique ground-truth spatial domains is unknown, we performed Louvain clustering with default resolution based on the latent embeddings learned by different methods. As shown in Fig. 2a and Supplementary Fig. S1a, PAST again achieved the overall best performance. Besides, PAST-E showed much more advantages than PAST-S in terms of various metrics, highlighting the benefit of external reference data.

Second, we qualitatively evaluate the performance of spatial visualization. Taking Slice 151673 as an example, we visualized the spatial domains characterized by clustering with a specified number of clusters and the ground-truth spatial domains on the spatial coordinates of ST data (Fig. 2b). The spatial domain assignment of the non-spatial approaches, namely Scanpy and Seurat, provided poor continuity and smoothness, which was consistent with the above quantitative results. SpaGCN, BayesSpace, CCST, DR-SC, and STAGATE could roughly follow the expected layer pattern in this slice, but the boundary of the identified domains was discontinuous with many outliers, which impaired the spatial clustering accuracy. PAST-S and PAST-E successfully characterized the expected cortical layer structures and achieved 6.3% and 10.9% improvement, respectively, compared to STAGATE in terms of AMI. We also visualized the spots using uniform manifold approximation and projection (UMAP) based on the latent embeddings learned by different methods (Fig. 2c). Expect for the white matter (WM), Scanpy and Seurat could hardly distinguish the cortical layers. SpaGCN, DR-SC, and STAGATE provided relatively better performance, while PAST successfully identified all the spatial structures and reconstructed the manifold of them. Besides, the spatial coordinate, UMAP and t-SNE visualization for the other 11 DLPFC sections also provided similar results (Supplementary Fig. S2-4).

Third, we examined the performance of latent embeddings for spatial trajectory inference. Based on the latent embeddings learned by different methods, we performed trajectory inference using PAGA on Slice 151673^29^ (Methods). As shown in Fig. 2d, PAST captured the well-known corticogenesis patterns^30, 31^ and inferred a nearly linear development trajectory from inner to outer layers, indicating the utility of PAST in revealing spatial development trajectory.

Fourth, we further examined the performance of latent embeddings for spatial pseudo-time analysis using DPT^32^ (Methods). As shown in Fig. 2e, PAST successfully characterized all the continuous development stages, while the baseline methods only captured the pseudo-time variation between WM and other cortical layers. Trajectory inference and pseudo-time analysis for other sections also provided similar results (Supplementary Fig. S5, 6), highlighting the potential of PAST to facilitate downstream analyses and biological implications.

### PAST takes full advantage of reference data constructed from various sources

The previous example on DLPFC sections utilized reference data constructed from the target section itself (PAST-S) and from the other sections (PAST-E). The prior information in external reference data is generally incomplete for the target ST data. Therefore, we further carefully examined the robustness of PAST-E to the completeness and scale of reference data, using STAGATE, the second-best method on DLPFC datasets, as the baseline.

To investigate the influence of incomplete reference data, we gradually left out spatial domains in external reference data, and obtained PAST-E-LO1, PAST-E-LO2, PAST-E-LO3, and PAST-E-LO4, which denote leaving out layer 1, layers 1 and 2, layers 1 through 3, and layers 1 through 4, respectively. The results shown in Fig. 3a and Supplementary Fig. S7a were in line with intuition since as the spatial domains are continuously left out, the effective prior information in the reference data is decreased. Except for PAST-E-LO4, the overall clustering performance of variants of PAST-E was superior to STAGATE (the horizontal dashed lines), and PAST-E performed well even based on the reference data containing only nearly half of shared domains, suggesting that PAST-E is robust to the completeness of reference data.

**Fig. 3.**
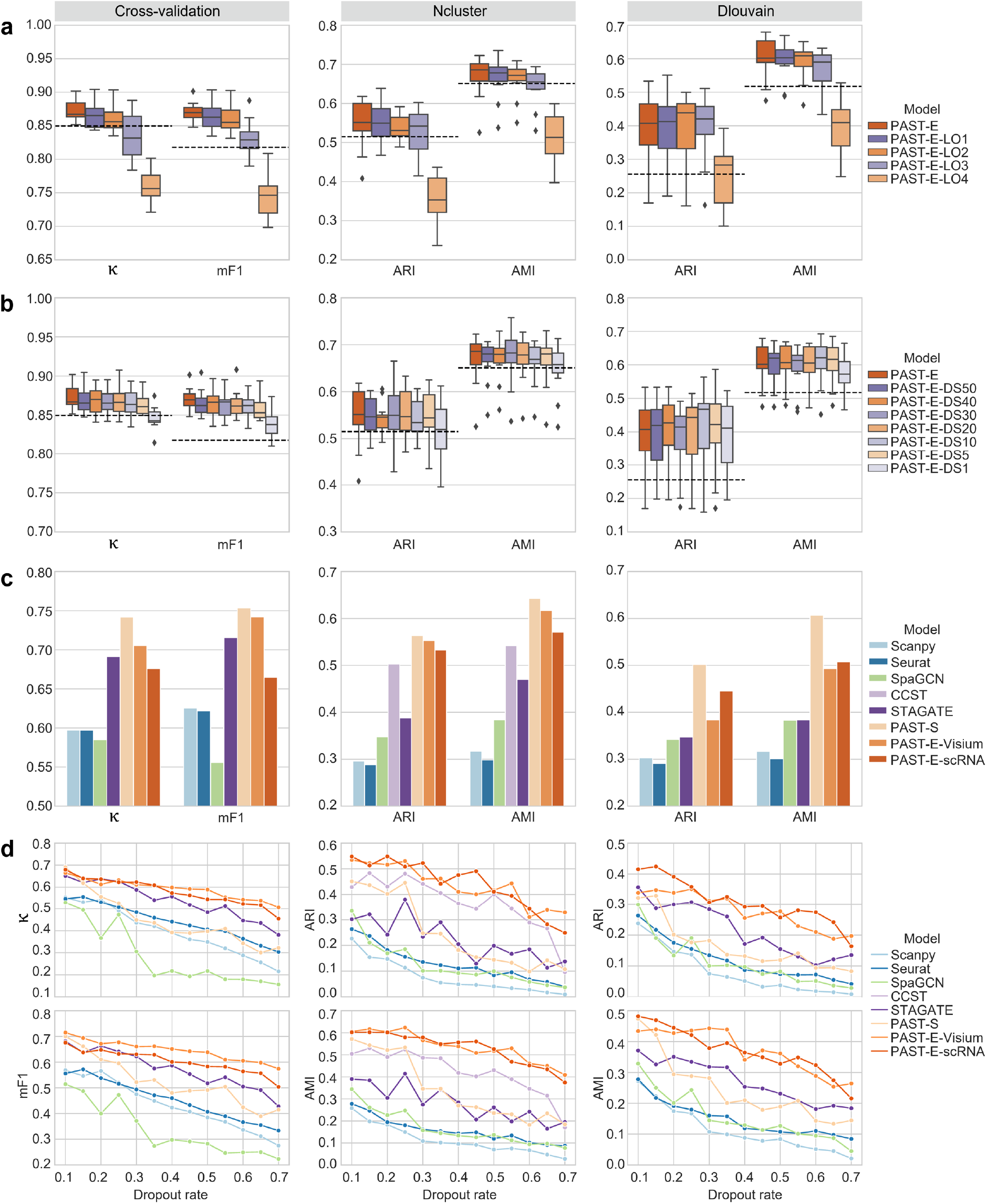
PAST takes full advantage of reference data constructed from various sources. Quantitative performance evaluation of spatial domain characterization on 12 DLPFC slices and STARmap MPVC dataset via supervised cross-validation and unsupervised spatial clustering with a specified number of clusters (Ncluster) and with default resolution (Dlouvain). The cross-validation performance was evaluated by the average score of Cohen’s kappa value (κ) and mean F1 score (mF1), respectively, in the 5-fold experiments. The spatial clustering performance was evaluated by adjusted rand index (ARI) and adjusted mutual information (AMI). The horizontal dashed lines represent the corresponding median scores of STAGATE. The center line, box limits and whiskers in the boxplots are the median, upper and lower quartiles, and 1.5× interquartile range, respectively. **a**, The spatial domains in external reference data of PAST were gradually left out. **b**, The spots in external reference data of PAST were downsampled to construct reference data with various scales. **c**, Performance evaluation of PAST on the STARmap MPVC dataset with reference data constructed from various sources. **d**, Performance evaluation of different methods under various dropout rates on MPVC.

To test the influence of limited number of spots in reference data, we randomly downsampled the spots in external reference data, and obtained PAST-E-DS50, PAST-E-DS40, PAST-E-DS30, PAST-E-DS20, PAST-E-DS10, PAST-E-DS5, and PAST-E-DS1, which denote remaining sampled 50%, 40%, 30%, 20%, 10%, 5%, and 1% of the spots, respectively. Compared with PAST-E which integrated prior information from more than 40,000 spots, the data scale of reference of PAST-E-DS50 through PAST-E-DS1 was only 21,949 through 425. As shown in Fig. 3b and Supplementary Fig. S7b, PAST-E with downsampled reference data consistently achieved satisfactory performance for supervised cross-validation and spatial clustering, until only 1% of the spots in reference data remained. We then performed two-sided Wilcoxon signed-rank tests on the quantitative results of different reference data scales. As shown in Supplementary Fig. S8, the scale of reference data, which may range from a few hundred to tens of thousands, has no significant impact on the performance of PAST-E, indicating that PAST-E takes full advantage of reference data and is robust to the data scale.

To further demonstrate that the reference data can be constructed from various sources in practice, we collected ST data from a STARmap dataset of the mouse primary visual cortex (MPVC)^33^. We constructed the reference data from the target ST data itself (referred to as PAST-S), a 10x Visium ST dataset of the mouse brain coronal^34^ (referred to as PAST-E-Visium), and a scRNA-seq dataset of the mouse cortex^35^ (referred to as PAST-E-scRNA). As shown in Fig. 3c and Supplementary Fig. S9a, PAST-S achieved significant improvement compared with the baseline methods for spatial domain characterization. The results of spatial coordinate, UMAP and t-SNE visualization, trajectory inference, and pseudo-time analysis also demonstrated the advantage of PAST-S (Supplementary Fig. S10). Contrary to the above example on DLPFC sections, PAST variants with external reference data (PAST-E-Visium and PAST-E-scRNA) performed worse than PAST-S, which may be due to the inconsistency of the tissues and protocols between target and reference data. Even that, PAST-E-Visium and PAST-E-scRNA still outperformed the baseline methods, indicating that PAST can make full use of reference data constructed from diverse biological contexts, and from other ST protocols or even scRNA-seq.

To mimic protocols that provide low coverage, we downsampled the reads in the target ST data by randomly dropping out the entries in the expression matrix to zero with a probability equal to the specified dropout rate. As shown in Fig. 3d and Supplementary Fig. S11, PAST variants with external reference data (PAST-E-Visium and PAST-E-scRNA) were less affected when the dropout rate increased, and consistently outperformed baseline methods when the dropout rate varies from 10% to 70%. Intuitively, the performance of PAST-S also deteriorated as the dropout rate increased, since the reference data, namely the target data itself, also became much noisier. Besides, we also showed that PAST is robust to model hyper-parameters including the number of neighbors and latent dimensions on the STARmap MPVC dataset (Supplementary Fig. S12, 13). Note that we did not evaluate the performance of BayesSpace on the STARmap MPVC dataset since it was specifically designed for ST and 10x Visium data^5^. Altogether, the results suggest the increased benefit of utilizing reference data when the target ST data has a higher degree of sparsity and noise.

### PAST capably captures local spatial patterns with self-attention mechanism

Next, we tested whether PAST could provide insights into local spatial patterns. We applied PAST to an osmFISH dataset profiled from the mouse somatosensory cortex (MSC)^36^, and utilized the target data itself as a reference. As shown in Fig. 4a and Supplementary Fig. S14, PAST-S effectively characterized spatial domains, and achieved significant improvement compared to the baseline methods in terms to quantitative evaluation of supervised cross-validation, and unsupervised spatial clustering with a specified number of clusters or with default resolution. Besides, PAST with different numbers of neighbors also demonstrated its consistent superiority (Supplementary Fig. S15). For spatial visualization, PAST-S successfully identified the expected structures and delineated the layer borders clearly, and the identified domains were more consistent with the original annotation of the dataset (Fig. 4b, c).

**Fig. 4.**
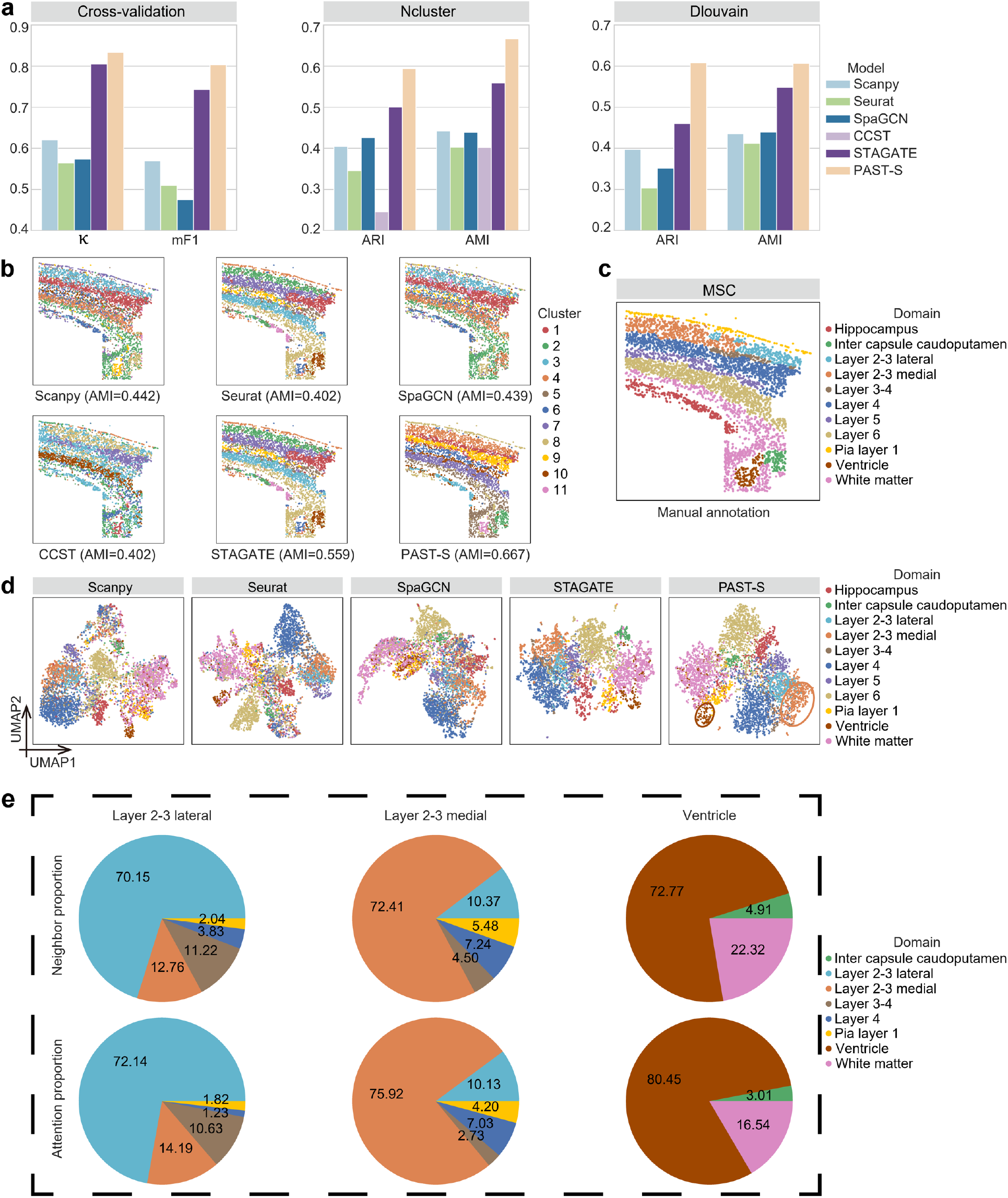
PAST capably captures local spatial patterns with self-attention mechanism. **a**, Quantitative performance evaluation of spatial domain characterization on the osmFISH MSC dataset via supervised cross-validation and unsupervised spatial clustering with a specified number of clusters (Ncluster) and with default resolution (Dlouvain). The cross-validation performance was evaluated by the average score of Cohen’s kappa value (κ) and mean F1 score (mF1), respectively, in the 5-fold experiments. The spatial clustering performance was evaluated by adjusted rand index (ARI) and adjusted mutual information (AMI). **b**, Visualization of the results of spatial clustering with a specified number of clusters on MSC. **c**, The manual annotation of MSC. **d**, UMAP visualization of spots in MSC. The blue, orange and brown circles denote Layer 2-3 lateral, Layer 2-3 medial and Ventricle, respectively. **e**, The neighbor proportion and attention proportion for border spots of domains Layer 2-3 lateral, Layer 2-3 medial and Ventricle, respectively.

We then explored the capability of the self-attention module of PAST-S in capturing local spatial patterns. As shown in Fig. 4d and Supplementary Fig. S16, compared with GCN-based methods like SpaGCN, PAST could effectively identify domains like Layer 2-3 lateral, Layer 2-3 medial and Ventricle in UMAP and t-SNE visualization (blue, orange and brown circles). For these three spatial domains, we selected the border spots whose spatial neighbors were originated from at least one other spatial domain, and calculated the neighbor proportion and attention proportion of each domain. More specifically, for each border spot of each domain, we recorded the proportion of different spatial domains in the 6 nearest neighboring spots, and calculated the average proportion for all the border spots of each domain, resulting in the neighbor proportion. In comparison with the neighbor proportion, the attention proportion recorded the attention-weighted proportion of different spatial domains in the 6 nearest neighboring spots. As shown in Fig. 4e, the self-attention mechanism successfully identified the neighboring spots from the same domain and improved the corresponding weights, i.e., from 70.15% to 72.14% for Layer 2-3 lateral, from 72.41% to 75.92% for Layer 2-3 medial and from 72.77% to 80.45% for Ventricle, indicating that self-attention mechanism enables border spots to aggregate more information from neighbor spots belonging to the same domain. Taken together, PAST could effectively capture the local spatial patterns with the self-attention mechanism and thus characterize the spatial correlation of the spots in ST data, especially the easily confused border spots.

### PAST enables scalable training and prediction on large data while preserving global spatial patterns

With the development of ST technologies, the number of profiled spots and genes increases exponentially, requiring the scalability of computational methods. Traditional mini-batch training strategy for deep learning is not suitable for the promising GNN because of the destruction to the connectivity and structure of graph^4, 22^, making GNN-based methods such as SpaGCN^6^ and CCST^7^ not applicable to large-scale ST data. STAGATE partitioned the profiled slice and constructed a graph for each part to enable training on large data, which may lead to the loss of global spatial patterns. We thus introduced a subgraph-based strategy for scalable training and prediction that considers both the randomness and connectivity of the graph and samples high-quality subgraphs based on a ripple walk sampler (RWS)^22^. RWS was originally proposed only for mini-batch training, and we further extended RWS from training to prediction by sampling subgraphs covering all spots of a dataset and aggregating the latent embeddings of different subgraphs in an ensemble manner (Methods).

We collected a Stereo-seq mouse olfactory bulb section (MOBS1) with 107,416 spots and 26,145 genes and analyzed the dataset on a standard desktop with an AMD Ryzen 7 Eight-Core 3800X CPU, 64GB of RAM and an NVIDIA GeForce RTX 2080 Ti GPU. We collected another Stereo-seq mouse olfactory bulb section (MOBS2) with 104,931 spots and 23,815 genes to construct the external reference for PAST. Note that we encountered memory errors when performing Seurat, SpaGCN and CCST on MOBS1. As shown in Fig. 5a and Supplementary Fig. S17, PAST-S and PAST-E, which incorporated self-prior information from MOBS1 itself and external-prior information from MOBS2, respectively, achieved the overall best performance for spatial domain characterization. PAST-S and PAST-E also achieved satisfactory performance in UMAP and t-SNE visualization (Supplementary Fig. S18). Besides, the spatial domains detected by PAST-S and PAST-E were spatially continuous and smooth, and the subependymal zone (SEZ) domain, denoted as Cluster 1, was only successfully characterized by these two variants of PAST (Fig. 5b, c). We performed differential expression gene (DEG) analysis for Cluster 1 of PAST-S and PAST-E, respectively. The results of both of these two variants showed that the top two DEGs were *Mbp* and *Plp1*, which are classical genes encoding myelin proteins^37^. Myelin proteins were usually used as markers for striatal contamination or staining to facilitate accurate dissection of the SEZ^37, 38^. We further performed gene ontology (GO) analysis for the top 10 DEGs with the lowest adjusted *p*-values of Cluster 1, and identified the most significant molecular function, i.e., structural constituent of myelin sheath, with false discovery rates (FDR) of 1.28e-04 and 9.33e-05 for PAST-S and PAST-E, respectively. The GO analysis results were well consistent with the molecular function of SEZ domain, i.e., deriving oligodendrocytes to form mature myelin sheaths around axons^39, 40^, suggesting the ability of PAST to facilitate functional implication.

**Fig. 5.**
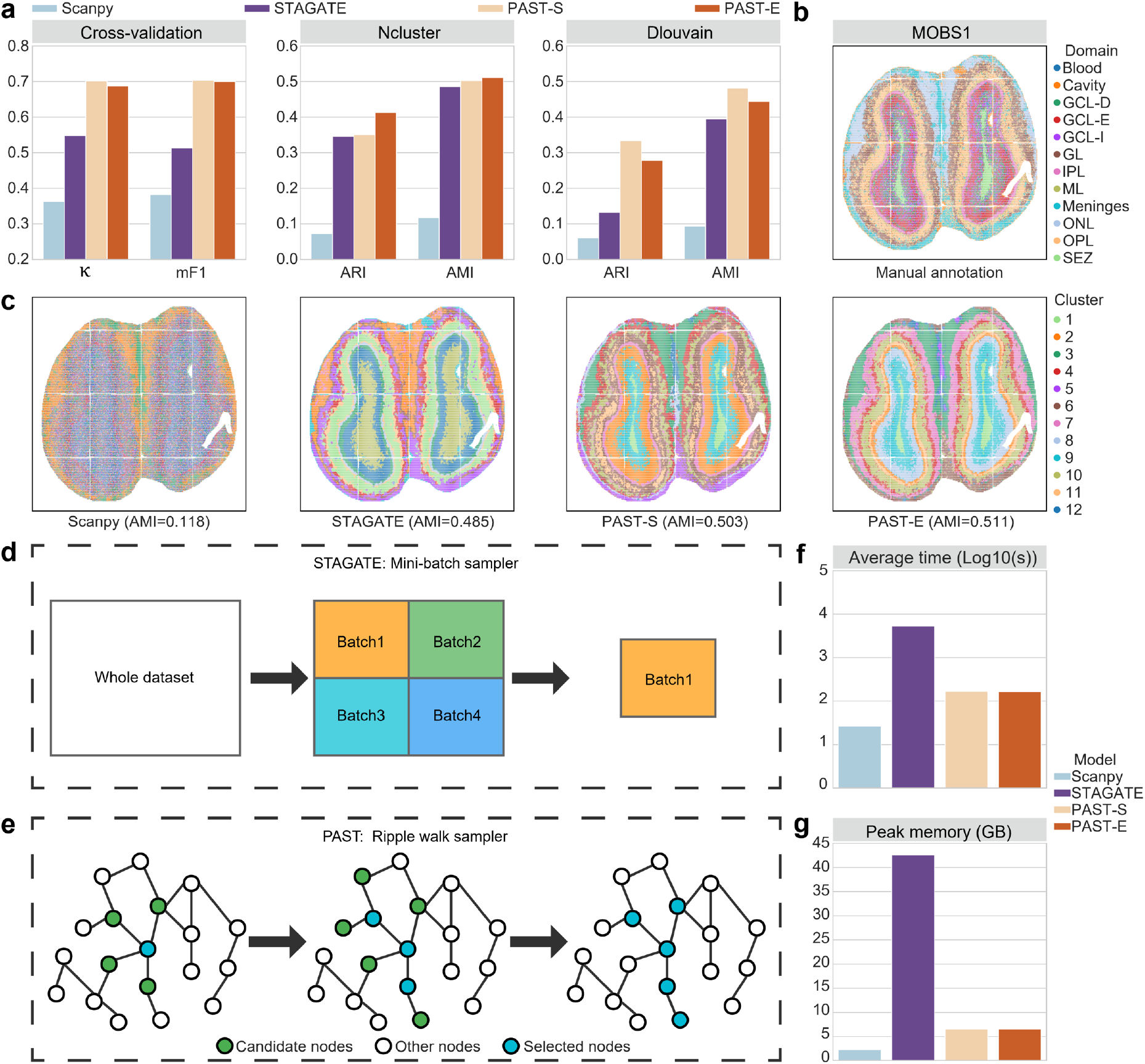
PAST enables scalable training and prediction on large data while preserving global spatial patterns. **a**, Quantitative performance evaluation of spatial domain characterization on the Stereo-seq MOBS1 dataset via supervised cross-validation and unsupervised spatial clustering with a specified number of clusters (Ncluster) and with default resolution (Dlouvain). The cross-validation performance was evaluated by the average score of Cohen’s kappa value (κ) and mean F1 score (mF1), respectively, in the 5-fold experiments. The spatial clustering performance was evaluated by adjusted rand index (ARI) and adjusted mutual information (AMI). **b**, The manual annotation of MOBS1. **c**, Visualization of the results of spatial clustering with a specified number of clusters on MOBS1. **d**, Sampling strategy of STAGATE for mini-batch training. **e**, Ripple walk sampling strategy of PAST for scalable subgraph-based training and prediction. **f**, Average time cost of different methods on MOBS1. **g**, Peak memory usage of different methods on MOBS1.

We also recorded the computational time and peak memory usage of different methods, especially the two GNN-based methods, i.e., STAGATE and PAST, which utilized different sampling strategies (Fig. 5d, e). As shown in Fig. 5f, the average computational time of PAST-S and PAST-E in 10 replicated experiments were approximately 30× less than STAGATE. In terms of peak memory usage, PAST-S and PAST-E also consumed significantly less memory than STAGATE (Fig. 5g), indicating that the ripple walk sampling strategy of PAST has a significant advantage over the mini-batch sampling strategy of STAGATE. Note that Scanpy also provided satisfactory scalability since its straightforward embedding strategy. Taken together, PAST enables scalable subgraph-based training and prediction while preserving global spatial patterns to better characterize spatial organizations.

### PAST facilitates automatic domain annotation and reveals biological implications

Computational methods for automatic spatial domain annotation are urgently needed given the exponential growth in the number of spots. However, to the best of our knowledge, there is no method proposed specifically for the automatic annotation of spatial domains in new ST data. We hence proposed a supervised annotation strategy based on the PAST framework and also applied this strategy to other spatial embedding methods (Methods). We first adopted the 12 DLPFC sections to quantitatively demonstrate the performance. Specifically, we used Slice 151673 to train PAST with reference constructed with the target dataset itself, and then obtained the joint embeddings of all the 12 sections with the trained PAST-S model. As shown in Fig. 6a, b and Supplementary Fig. S19, UMAP and t-SNE visualization based on the embeddings of spots in all the 12 sections illustrated the capability of PAST in batch correction and domain identification. We next utilized the embeddings and domain labels of Slice 151673 to train a support vector machine classifier and annotated the spatial domains in other sections. For quantitative comparison, PAST consistently outperformed the baseline methods (Fig. 6c), and achieved significantly higher scores of Acc, κ, mF1 and wF1 than STAGATE, the second-best method, with *p*-values of 0.0034, 0.0024, 0.0005, and 0.0005, respectively, in one-sided paired Wilcoxon signed-rank tests.

**Fig. 6.**
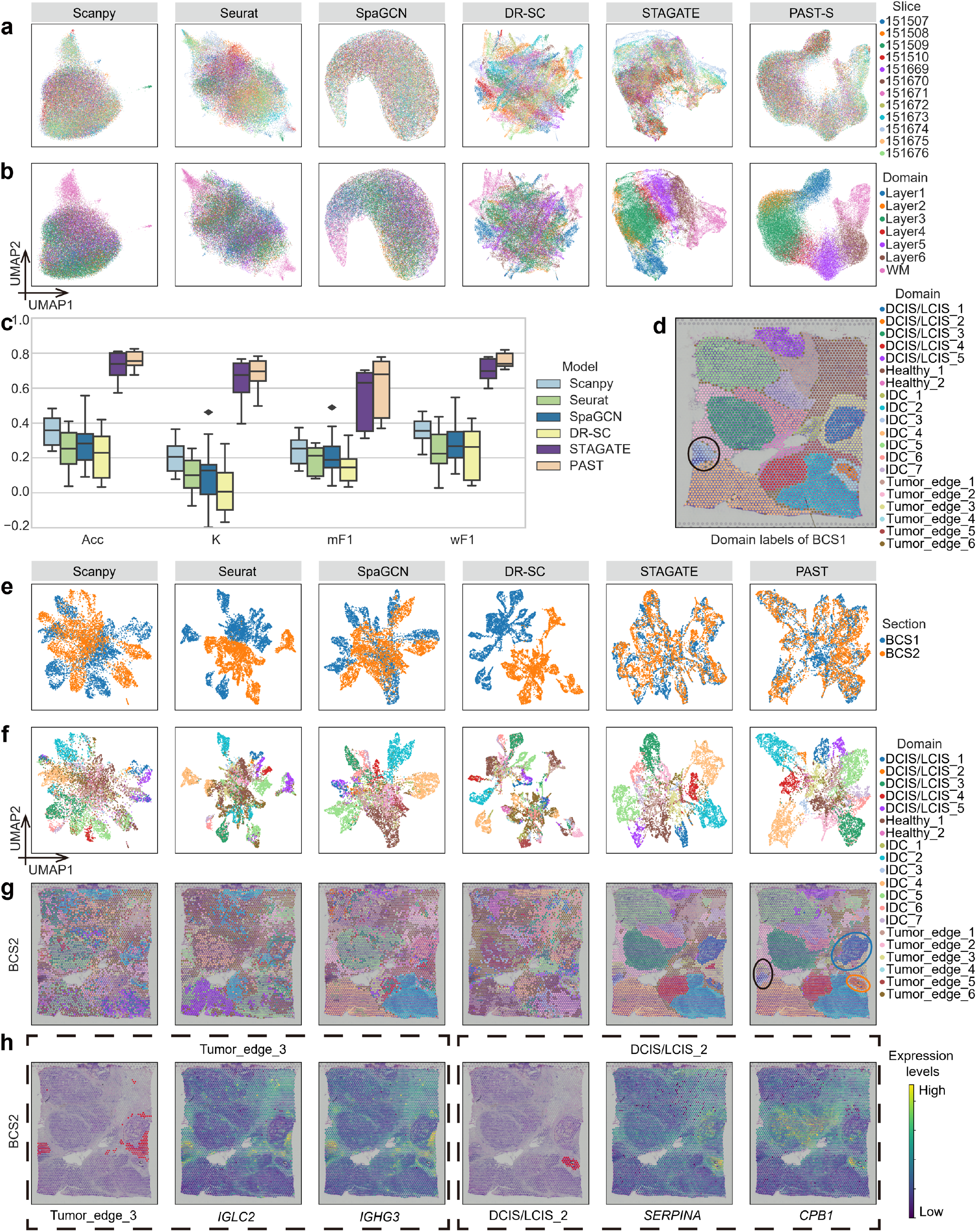
PAST facilitates automatic domain annotation and reveals biological implications. UMAP visualization of spots in all the 12 DLPFC slices colored by **a**, slices and **b**, spatial domains. **c**, Performance of supervised annotation evaluated by accuracy (Acc), Cohen’s kappa value (κ), mean F1 score (mF1) and weighted F1 score (wF1) where Slice 151673 was taken as the training set and other 11 slices were taken as test sets. The center line, box limits and whiskers in the boxplots are the median, upper and lower quartiles, and 1.5× interquartile range, respectively. **d**, The manual annotation of BCS1, in which the IDC_3 tumor region was black circled. UMAP visualization spots in BCS1/2 sections colored by **e**, sections and **f**, spatial domains. **g**, The supervised annotation results on BCS2 by different methods. The black circle denotes the supervised annotated IDC_3 and Tumor_edge_3, while the blue and orange circles denote the supervised annotated DCIS/LCIS_1 and DCIS/LCIS_2, respectively. **h**, The annotation results of Tumor_edge_3 and DCIS/LCIS_2 by PAST and the spatial expression patterns of BCS1-marker genes on BCS2.

We further collected two 10x Visium datasets of the human breast cancer sections (BCSs). Similarly, we utilized BCS1 (Fig. 6d) to train PAST-S and obtained the joint embedding of the two sections. PAST again successfully corrected the batch effect between the sections and separated spots from different domains (Fig. 6e, f and Supplementary Fig. S20). We next used the first section (BCS1) which comes with spatial domain annotations to train a classifier, and annotated the domains in BCS2, another section that comes without spatial domain annotations (Fig. 6g). For the evaluation of annotation results, we identified marker genes for each domain based on the ground-truth labels on BCS1 (referred to as BCS1-marker genes), and found that BCS1-marker genes of each domain were also differentially expressed in most of the corresponding PAST-annotated tumor areas on BCS2 (Supplementary Fig. S21), indicating the ability of PAST to reveal biological variation among tumor areas. In addition, PAST successfully captured BCS2-specific spatial domains by separating the IDC_3 region on BCS1 into IDC_3 and Tumor_edge_3 on BCS2 (black circles in Fig. 6d, g). The BCS1-marker genes of Tumor_edge_3 also demonstrated the unique spatial pattern of the newly annotated Tumor_edge_3 on BCS2, which further suggsets the accuracy of annotation results on BCS2 (Fig. 6h).

We also performed differential expression gene analysis for Tumor_edge_3 region based on the annotation results of PAST on BCS2, and identified the five most significant differential expression genes (DEGs) including *IGHG3, IGKC, IGLC2, IGHG4* and *IGHG1* with adjusted *p*-values of 1.11e-83, 1.86e-78, 9.18e-74, 9.18e-74 and 1.66e-69, respectively. According to previous literature, the linkage of *IGHG1/3/4* provides information for prognosis of diseases like autoimmunity and malignancy^41^, *IGKC*, serving as a robust immune marker, could predict metastasis-free survival and response to chemotherapy^42^, and *IGLC2* was proved to be a significant prognostic biomarker for triple-negative breast cancer patients^43^. The gene ontology (GO) enrichment for the top 10 DEGs of Tumor_edge_3 also significantly revealed lots of biological immune processes (Supplementary Fig. S22).

PAST also successfully captured the biological difference between DCIS/LCIS_1 and DCIS/LCIS_2 (blue and orange circles in Fig. 6g) and accurately annotated DCIS/LCIS_2 on BCS2, which was missed by other methods. The spatial expression visualization of BCS1-marker genes of DCIS/LCIS_2 on BCS2 illustrated the accurate and superior annotation of DCIS/LCIS_2 by PAST (Fig. 6h). To summarize, PAST can not only effectively facilitate automatic spatial domain annotation, but also identify test set-specific spatial patterns and reveal the corresponding biological implications.

## Discussion

Rapid development of ST sequencing technologies has increased the demand for effective characterization of spatial domains. In this article, we introduce PAST, a comprehensive and scalable framework that effectively integrates prior information via Bayesian neural networks and models spatial correlations based on variational graph convolutional auto-encoders with self-attention mechanism and ripple walk sampler strategy. With comprehensive experiments on multiple datasets generated by various technologies, we have shown the superior information fusion capabilities of PAST to take full advantage of spatial locations and reference data from various sources. We have also demonstrated the successful application of PAST in downstream tasks including spatial domain characterization, ST visualization, spatial trajectory inference, and pseudo-time analysis. Besides, PAST not only enables automatic annotation of spatial domains for newly sequenced ST data, but also provides biological insights into the annotated domains.

Certainly, there are several avenues for improving PAST. First, as a generative model, we can explore the application of PAST in ST data simulation by adding more complex assumptions on latent distribution^44^. Second, the rapid development of GNN field also opens new horizons for better characterization of spatial patterns. Finally, as a representation learning model, PAST could be extended to characterize spatial domains in other spatial omics data, such as spatial-ATAC-seq data^45^.

## Methods

### The model of PAST

PAST is a weakly supervised representation learning framework built upon variational graph convolutional auto-encoder^46^ and it takes three kinds of data as input, including preprocessed target gene expression matrix, target spatial coordinate matrix and prior gene expression matrix (Fig. 1).

In the forward propagation phase, we first construct a *k*-NN graph using the target spatial coordinates with default parameter *k* = 6 and sample high-quality subgraphs based on the *k*-NN graph using a ripple walk sampler. We then feed the subgraphs and the target and prior gene expression matrices into the encoder of PAST, where the prior gene expression matrix provides reference information for the Bayesian neural network (BNN) module. Based on the output of encoder, we utilize the reparameterization trick^46^ to obtain multivariate normally distributed samples and then feed the samples into the decoder to get a reconstructed target gene expression matrix. In the backward propagation phase, we calculate the gradient of each parameter in the model according to the loss function and then update parameters using the Adam optimizer with default initial learning rate *lr* = 0.001. The detailed model structure and hyper-parameter settings of PAST are summarized in Supplementary Table S1.

#### Encoder

The encoder of PAST is a three-layer neural network. The first layer consists of a BNN module and a parallel fully connected module, and the output of the first layer is the concatenation of the two modules. The second and third layers are two stacked self-attention modules. The input of self-attention module includes a feature matrix from the last layer and the *k*-NN graph constructed with spatial coordinates of the target data.

#### Gaussian distribution assumption

As suggested by variational auto-encoder (VAE), we assume the prior of latent embeddings of spots is centered isotropic multivariate Gaussian distribution, use two fully connected layers to obtain the mean vector and logarithmic variance vector of the latent space based on the output of encoder, and obtain multivariate normally distributed samples through the reparameterization trick^46^.

#### Decoder

The decoder of PAST is also a three-layer network, where the first and second layers are stacked self-attention modules and the third layer is a fully connected layer. The decoder takes multivariate normally distributed samples from the latent Gaussian sampler as input and then outputs the reconstructed gene expression vectors of the target data.

### Bayesian neural network

Bayesian neural network (BNN) introduces uncertainty into weights of the neural network to improve generalization in non-linear regression problems^47^. Here we use BNN to integrate prior information from reference datasets.

Parameters of the BNN module could be represented as a *d*_*in*_ × *d*_*out*_ matrix ***P***, and each element in ***P*** is subject to a normal distribution,

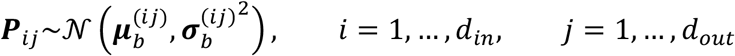

where 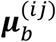 and 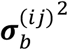 are elements in the *i*-th row and *j*-th column of the mean matrix ***μ*** and variance matrix 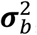, respectively. We get normally distributed parameters ***P***_*ij*_ through reparameterization trick^46^ in each forward propagation process. Given an input matrix ***M*** of shape *B* × *d*_*in*_ in a mini-batch, the output matrix of shape *B* × *d*_*out*_ of BNN module is

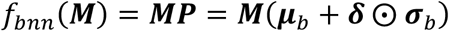

where *f*_*bnn*_(·) denotes the function of BNN module, ***δ*** is the standard gaussian noise matrix and ⨀ denotes element-wise product.

To further integrate prior information in existing data from various technologies, we assume elements in ***P*** are subject to a prior gaussian distribution parameterized by ***μ***_*p*_ and 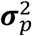, and use Kullback-Leibler divergence to restrict the distance between the BNN parameter distribution and the prior distribution^47^,

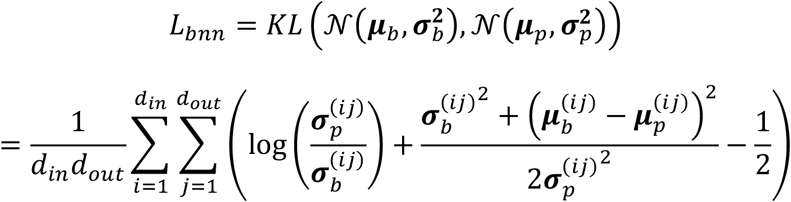

where ***μ***_*p*_ is the principle component analysis (PCA) projection weight matrix of the prior gene expression matrix and ***σ***_*p*_ contains elements equal to one-tenth standard deviation of ***μ***_*p*_.

### Construction of prior gene expression matrix

PAST can integrate the prior information contained in reference data derived from various sources.

First, we consider the situation where external ST data of the same tissue and from the same protocol with the target ST data is available. Platforms like 10x Visium, Stereo-seq and Slide-seq tend to produce multiple sequencing samples of the same tissue once a time^16-18^, and these samples from the same platform can be used as reference data for each other.

Second, we consider the situation where external ST data of similar tissue and from other protocols is accessible. Recently, various ST sequencing technologies, including 10x Visium, STARmap, osmFISH, Slide-seq and Stereo-seq, have produced numerous ST datasets^16-20, 36^, and all the datasets from different platforms can serve as potential references to each other.

Third, external scRNA-seq data from similar tissue can also be utilized to construct a prior gene expression matrix for target ST data. Since the development of single-cell technology has generated an enormous amount of scRNA-seq data, we believe that integrating scRNA-seq data to analyze ST data will dramatically expand the applicability of ST data.

Fourth, when there is not any relevant external data can be accessed, the target data itself can also be used as a reference in this circumstance, and we have demonstrated that the self-prior information can also effectively promote the performance for deciphering spatial domains.

After obtaining the reference gene expression matrix from various sources, we first align the gene set of the reference dataset to that of the target ST dataset, and then apply the same preprocessing procedure as the target dataset to the reference dataset, ensuring that the reference and the target dataset share a consistent feature space. If there are spatial domain labels accompanying the reference dataset, we construct a pseudo-bulk gene expression matrix based on the domain labels, the insight of which is that bulk data is more robust to technological noise than single-cell data^48^ and pseudo-bulk data constructed with annotation labels contains more valuable category information^14^. The step-by-step construction of pseudo-bulk gene expression matrix is described in Algorithm 1 of Supplementary Notes. If there are no spatial domain labels accompanying the reference dataset, we directly utilize the processed reference dataset as the prior gene expression matrix. Finally, we perform PCA on the prior gene expression matrix and use the PCA projection weight matrix as the means in the BNN module.

### Self-attention mechanism

Self-attention mechanism is the key mechanism of transformer for capturing the relationship between words in a given sentence in machine translation tasks, showing superior performance compared with other recurrent neural network-based models^21^. Recent advances have revealed that gene expression levels of neighboring spots in a tissue microenvironment tend to be correlated^4, 8^. Inspired by the above understanding, PAST adopts a self-attention mechanism to model the relationship between neighboring spots (Fig. 1).

The input of the self-attention module is a feature matrix ***M*** and the corresponding *k*-NN graph 𝒢. 𝒢 is constructed using spatial coordinates with *k* = 6. The input matrix ***M*** is mapped to obtain query ***Q***, key ***K*** and value ***V*** through three fully connected neural networks, and the adjacent matrix 𝒜 can then be calculated as follows,

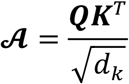

where *d*_*k*_ is the feature dimension of query ***Q*** and key ***K***. The attention weight matrix ***A*** can be obtained by normalizing the adjacent matrix 𝒜

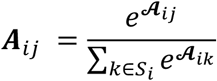

where ***A***_*ij*_ is the attention weight between spots *i* and *j*, and *S*_*i*_ denotes the set of neighbors of spot *i* in 𝒢. To avoid excessive spatial smoothing of hidden features, we add a shortcut in the self-attention module^21^ and the output of self-attention module can be written as,

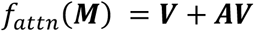

where *f*_*attn*_(·) denotes the function of self-attention module.

### Loss function

Analysis of omics data is susceptible to sequencing noise such as dropout^49^, and recent studies have proved that it is beneficial to consider data sparsity and dropout in spatial transcriptomic data^15^. Here we design sparsity adaptive reconstruction loss based on the insight that the higher the data sparsity, the more serious the dropout phenomenon is, and the higher probability the zero gene expression observations are false negative. Given a mini-batch containing *B* spots and *M* genes, the sparsity adaptive reconstruction loss is shown as,

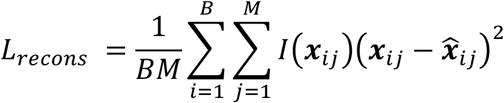

where ***x***_*ij*_ denotes the profiled expression level of gene *j* in spot *i*, 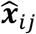 denotes the reconstructed expression level of gene *j* in spot *i*, and the function ***I***(·) is

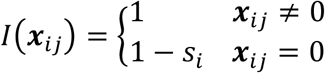

where *s*_*i*_ represents the data sparsity of spot *i* across all genes.

We assume the low-dimensional embeddings follow multivariate Gaussian distribution and the prior of latent embeddings is the centered isotropic multivariate Gaussian distribution. The distance between the latent distribution and the prior distribution is restricted by Kullback-

Leibler divergence^46^:

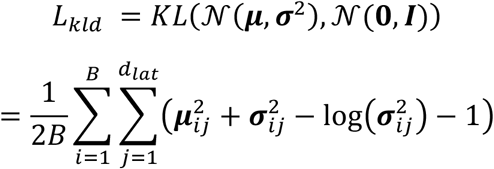

where ***μ*** and ***σ*** are the mean vectors and standard deviation vectors of latent space, respectively, while *d*_*lat*_ denotes the dimension of latent space.

Metric learning has been proposed to model the distance between samples, making the distance between positive sample pairs closer and the distance between negative sample pairs farther^25^. Samples in spatial transcriptomics are of strong spatial autocorrelation, which means that spatial neighboring spots may have similar gene expression patterns and tend to belong to the same functional structure^4^. Here we design a metric learning loss function to restrict the distance of low-dimensional representations between neighbor samples,

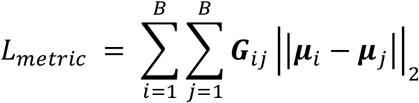

where ***G*** is the weighted spatial neighbor graph and the connection weights between samples are calculated using the student’s t distribution kernel,

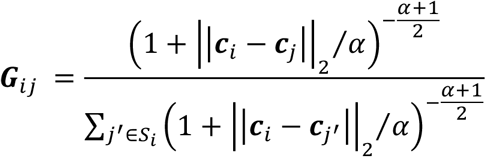

where ***c***_*i*_ denotes the spatial coordinate of spot *i, S*_*i*_ denotes the set of neighbors of spot *i, α* is a scale factor that is set to 1 by default.

As described in the above section, we integrate prior information through the Bayesian neural network^47^ module and restrict the distance between parameter of the BNN module and prior distribution through *L*_*bnn*_ loss function.

The objective of PAST is to minimize the combination of the above four parts of loss function:

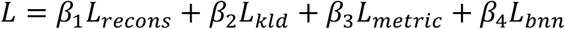

where *β*_1_, *β*_2_, *β*_3_, *β*_4_ are coefficients served as a trade-off between the four parts and we set *β*_1_ = 1, *β*_2_ = 1, *β*_3_ = 1, *β*_4_ = 1 by default.

### Ripple walk sampler

Since the damage to nodes connectivity and overall graph structure limits the usage of traditional mini-batch random sampling strategy, neighbors explosion and node dependence are common problems when training graph neural networks on large graph-structured datasets^22^. In order to extend our model to large-scale application scenarios, we adopt a subgraph-based sampling strategy called ripple walk sampler, which samples high-quality subgraphs to constitute mini-batch training and prediction under the premise of satisfying randomness and connectivity^22^, shown as Algorithm 2 in Supplementary Notes.

#### Ripple walk training

Ripple walk training (RWT) was proposed based on a ripple walk sampler for scalable mini-batch training. Following the settings in original paper^22^, we set the parameter expansion ratio *r* = 0.5 and adjust the number of subgraphs within one training iteration *S* to satisfy *BS* = 10, given a user-specified batch size *B*. The detailed training process of PAST with ripple walk sampler strategy is described in Algorithm 3 of Supplementary Notes.

#### Ripple walk prediction

Ripple walk sampler, which was proposed only for ripple walk training ^22^, could not guarantee full coverage of all samples. Here we refine the ripple walk sampler by selecting initial nodes from the set of non-sampled nodes so that we could obtain a set of subgraphs covering all spots in ST datasets, thus allowing subgraph-based prediction on large datasets. We also propose an ensemble strategy to get final latent embeddings. The proposed step-by-step ripple walk prediction procedure is shown in Algorithm 4 of Supplementary Notes.

### Automatic annotation

We propose a strategy to utilize PAST for the automatic annotation of spatial domains across multiple datasets in a supervised manner. Considering a dataset with spatial domain labels, which is denoted as anchor dataset, and multiple unannotated datasets, we propose to obtain joint latent embedding of all the datasets and then predict the spatial domain of the unannotated datasets with a support vector machine (SVM) trained on the anchor dataset.

Specifically, in the joint embedding step, we first train PAST on the anchor dataset with a pseudo-bulk prior matrix constructed with the anchor dataset itself based on the domain labels, and then apply the trained PAST on the anchor dataset and other unannotated datasets to obtain joint embeddings. In the prediction step, we train an SVM with default parameters on the anchor dataset, as suggested by two benchmark studies on supervised cell type identification for single-cell RNA-seq data^27, 28^. We then annotate the spatial domain of spots in other datasets using the trained classifier.

### Data collection

We collected spatial transcriptomic (ST) datasets generated by different technologies, including 10x Visium, STARmap, osmFISH, and Stereo-seq, for the performance evaluation of PAST. A summary of the ST datasets and the corresponding reference data is provided in Supplementary Table S2.

The dorsolateral prefrontal cortex (DLPFC) dataset contains 12 sections generated by 10x Visium from 3 independent neurotypical adult donors. The layer labels (Layer1-6, WM) of the 12 DLPFC sections were manually annotated based on cytoarchitecture and selected gene markers in a supervised manner^26^. The STARmap mouse primary visual cortex (MPVC) dataset contains the spatial location and gene expression of 1207 spots and is composed of 7 spatial domains^33^. A 10x Visium mouse brain coronal dataset containing 15 regions^34^ and a scRNA-seq mouse primary visual cortex dataset containing 23 regions^35^ were also collected as external reference datasets for better characterization of the target STARmap MPVC dataset. The osmFISH mouse somatosensory cortex (MSC) dataset, containing expression levels of 6471 cells on 33 target genes, was separated into 11 spatial domains in the original study^36^. The Stereo-seq mouse olfactory bulb (MOBS1) dataset, which measured spatial location and genome-wide gene expression levels of 107,416 spots, was manually separated into 12 spatial domains^50^. We also collected another mouse olfactory bulb (MOBS2) dataset generated by Stereo-seq containing 11 regions as the reference of MOBS1^50^. Two human breast cancer datasets (BCS1 and BCS2) were collected from the 10x Genomics platform for the supervised annotation experiment, where BCS1 served as the training set while BCS2 was annotated with the trained classifier. The domain labels of BCS1 were obtained from SEDR^51^.

### Data preprocessing

Raw target gene expression matrix of unique molecular identifier (UMI) count is stored in a *N* × *M* matrix, where *N* denotes the number of spots and *M* denotes the number of genes. Following the data preprocessing tutorial for spatial transcriptomic data implemented in Scanpy package^10^, we filtered the genes expressed in fewer than three spots, divided the gene expression level of each spot by total UMI counts across all genes, multiplied them by the median of total counts for spots, and then transformed the normalized matrix into a log scale.

We further selected genes that showed high-level spatial autocorrelation by calculating Geary’s *C*^52^ index of each gene, which was used to evaluate the spatial correlation of selected spatially variable genes in recent studies^6, 53^. For the calculation of Geary’s *C* index, we first construct a *k* -NN graph using the spatial coordinate matrix of target data with default parameter *k* = 30. Let ***w***_*ij*_ denote the element of *k*-NN graph in the *i*-th row and *j*-th column, which is equal to 1 if spots *i* and *j* are neighbors and 0 otherwise. Then given a gene expression vector ***x*** across *N* spots, the Geary’s *C* index of each gene can be calculated as follows,

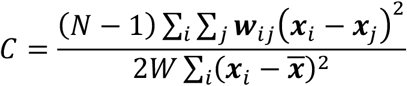

where *W* is the sum of all ***w***_*ij*_, ***x***_*i*_ denotes the gene expression in the *i*-th spot and ***x*** is the average expression value across all spots. To get an index that is positively related to spatial autocorrelation, we calculated *C*^*^ = 1 − *C* as transformed Geary’s *C* index. By default, we selected 3000 spatial variable genes with the highest transformed Geary’s *C* index in the preprocessing procedure.

### Baseline methods

The performance of PAST is compared with seven baseline methods with default parameters suggested in their tutorials for ST data analysis, including Scanpy^10^, Seurat^11^, SpaGCN^6^, BayesSpace^5^, CCST^7^, DR-SC^9^ and STAGATE^8^. The quantitative evaluation of PAST and all the baseline methods on all the datasets is summarized in Supplementary Table S3-S6.

Specifically, Scanpy, Seurat, SpaGCN, DR-SC, STAGATE could provide latent embeddings of spots for downstream analysis like visualization and clustering, while BayesSpace and CCST are spatial clustering methods that do not provide latent features.

In terms of applicable technology platforms, BayesSpace was specially designed for ST and 10x Visium, while DR-SC could be applied to several platforms, including 10x Visium, Spatial Transcriptomics, seqFISH, merFISH and Slide-seqv2. Other baseline methods are not constrained by the technology platforms. Therefore, BayesSpace was only applied to 10x Visium DLPFC datasets for spatial clustering, while DR-SC was used for spatial clustering on 10x Visium DLPFC datasets and automatic annotation on 10x Visium BCS1/2 datasets. Due to the large data scale, only Scanpy, STAGATE and PAST can be applied to the Stereo-seq MOBS1 dataset.

In terms of spatial embedding, we set the latent dimensions of all methods according to the suggestion of corresponding original study or tutorial, i.e., 15 for BayesSpace and DR-SC, 30 for STAGATE, 50 for other methods including Scanpy, Seurat, SpaGCN and PAST, in all experiments except for the osmFISH MSC dataset, since there are only 33 target genes measured in this dataset. On the osmFISH MSC dataset, we set the latent dimensions of Scanpy, Seurat, SpaGCN and PAST to 30 instead of 50 and keep default latent dimension settings for STAGATE.

Besides, to ensure a fair comparison of time and memory cost and performance, we set the training batch size of STAGATE and PAST uniformly to 3600 on the large-scale MOBS1, which is suggested by the tutorial of batch training strategy in STAGATE.

### Visualization

In all experiments, we obtained the low-dimensional representation with PAST and other spatial embedding methods, and then reduced the dimension to two using UMAP^54^ and t-SNE^55^ algorithms for visualization. UMAP and t-SNE algorithm has already been implemented in Scanpy package and the two-dimensional data were visualized via functions of *scanpy*.*pl*.*umap()* and *scanpy*.*pl*.*tsne()*.

### Clustering

For all the ST datasets, we considered two scenarios for spatial clustering based on the embeddings obtained by PAST and other methods. In the first scenario, we assumed the number of spatial domains is already known and utilized the clustering method suggested in the original paper or tutorial (Ncluster). Specifically, for Scanpy^10^ and Seurat^11^, we implemented a binary search strategy to tune the resolution of their default clustering method (Leiden and Louvain algorithm, respectively) to make the number of clusters and the number of spatial domains as close as possible. For SpaGCN, BayesSpace, CCST and DR-SC which could inherently output clustering results given a specified number of clusters, we set the number of clusters to be the number of unique spatial domains. For STAGATE and PAST, we performed the mclust algorithm which obtain clusters using gaussian finite mixture models with default parameters^56^.

In the second scenario, we assumed the number of spatial domains is not known for researchers because of the lack of knowledge about the corresponding organ or tissue. This scenario is more general than the first one. We adopted Louvain, one of the most commonly used community-based clustering algorithms that do not need a specified number of clusters, to perform spatial clustering (Dlouvain) based on latent embeddings obtained by all dimension reduction methods, including Scanpy, Seurat, SpaGCN, DR-SC, STAGATE and PAST. The resolution parameter of Louvain algorithm is set to 1 by default as suggested in the Scanpy tutorial.

### Trajectory inference

As suggested by STAGATE^8^, the PAGA algorithm^29^ was performed on embeddings of different methods based on the spatial domain labels to infer the developmental trajectories of different functional regions. PAGA algorithm was already implemented in Scanpy package and we visualized the spatial trajectory by the *scanpy*.*pl*.*paga()* function.

### Pseudo-time analysis

We used diffusion pseudotime (DPT) to estimate the temporal order of different spots based on their embeddings^32^. The Scanpy package also implemented the DPT algorithm and we performed DPT analysis on embeddings of each method by the functions of *scanpy*.*ti*.*diffmap()* and *scanpy*.*tl*.*dpt()*.

### Evaluation of embedding and annotation

We evaluated the information contained in the embeddings for predicting the true spatial domain labels through cross-validation. Following benchmark studies on supervised cell type identification for scRNA-seq data^28^, we adopted the support vector machine (SVM) with default parameters as the classifier. Specifically, we split all spots into five parts and conducted 5-fold cross-validation experiments, iteratively predicting spatial domain labels of the spots in each fold with the model trained with the remaining four folds.

Accuracy (Acc), Cohen’s kappa value (*κ*), mean F1 score (mF1) and weighted F1 score (wF1) were calculated to evaluate the prediction results of classifier in cross-validation and automatic annotation experiment. Accuracy score is the common metric for supervised prediction, and is calculated as follows:

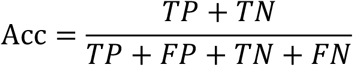

where *TP, FP, TN, FN* represent true positive, false positive, true negative and false negative, respectively. Cohen’s kappa value reflects the agreements between inter-annotators on a classification problem^57^, and is calculated as follows:

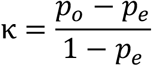

where *p*_*o*_ is the observed agreement ratio, *p*_*e*_ is the expected agreement when inter-annotators assign labels randomly. F1 score is the harmonic mean of precision score and recall score of binary classification problems^58^, which could be calculated as follows:

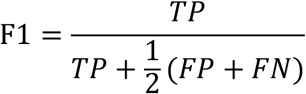

In multi-classification problems, we treated samples from a given category as positive and others as negative, calculated F1 score for each category and then obtained mF1 by calculating the mean F1 score of all categories. Besides, considering the label imbalance problem, we also calculated the weighted mean F1 score, denoted as wF1, where the proportions of different categories were taken as weights.

### Evaluation of clustering

To measure the agreement between clustering results and ground truth labels, we adopted six common clustering metrics, including adjusted rand index (ARI), adjusted mutual information (AMI), normalized mutual information (NMI), fowlkes-mallows index (FMI), completeness (Comp) and homogeneity (Homo).

Rand index denotes the probability that the clustering result and the true annotation will agree on a randomly selected set of samples. After consistent adjustment of the expectations of rand index, we can obtain ARI which ranges from 0 to 1 as follows:

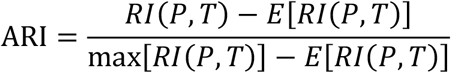

where *RI*(·) denotes the function of rand index, *T* denotes the true labels of samples, and *P* denotes the clustering result.

Both AMI and NMI originate from Mutual Information (MI), which measures the similarity between the clustering result and true labels. AMI, ranging from -1 to 1, is calculated as follows:

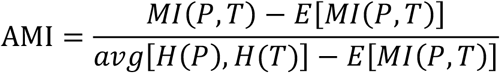

where *MI*(·) denotes the function of mutual information, *E*(·) is the expectation function and *H*(·) denotes the function of information entropy. NMI also ranges from 0 to 1, and can be calculated as follows:

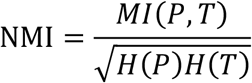

FMI is defined as the geometric mean of pairwise precision and recall. FMI also ranges from 0 to 1, and is calculated as follows:

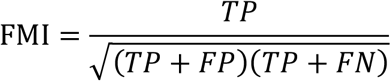

where *TP* (True Positive) is the number of pair of samples belonging to the same cluster in both the true labels and predicted labels, FP (False Positive) is defined as the number of pair of spots belonging to the same cluster in the true labels but not in the predicted labels, and FN (False Negative) is the number of pair of points belonging to the same cluster in the predicted labels but not in true labels.

Completeness means that all members of a given class are assigned to the same cluster, while homogeneity is defined as each cluster containing only members of a single class.

Bounded below by 0 and above by 1, completeness and homogeneity are calculated as follows:

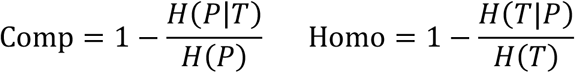

## Data availability

All target data and corresponding external prior data are available from original studies. The 10x Visium dorsolateral prefrontal cortex (DLPFC)^26^ data is accessible through the spatialLIBD R package http://spatial.libd.org/spatialLIBD. The STARmap mouse primary visual cortex (MPVC) data^33^ is available at https://www.starmapresources.org/data. Two reference datasets, i.e., 10x Visium mouse brain coronal dataset and scRNA-seq mouse primary visual cortex dataset can be downloaded by the squidpy package^34^ and from dropbox https://www.dropbox.com/s/cuowvm4vrf65pvq/allen_cortex.rds?dl=1, respectively. The osmFISH mouse somatosensory cortex (MSC) dataset can be obtained from http://linnarssonlab.org/osmFISH/availability/. The Stereo-seq mouse olfactory bulb sections 1 (MOBS1) and 2 (MOBS2) are accessible at https://db.cngb.org/stomics/mosta/download.html. The 10x Visium human breast cancer sections 1 (BCS1) and 2 (BCS2) can be downloaded from 10x Genomics platform https://www.10xgenomics.com/resources/datasets/human-breast-cancer-block-a-section-1-1-standard-1-1-0 and https://www.10xgenomics.com/resources/datasets/human-breast-cancer-block-a-section-2-1-standard-1-1-0, respectively.

## Code availability

The PAST algorithm is implemented in Python and the source code for reproduction is available at Github (https://github.com/lizhen18THU/PAST). We also provided detailed documentation and step-by-step tutorials for applying PAST to ST data generated by different technologies at Read the Docs website (https://past.readthedocs.io/en/latest).

## Acknowledgements

This work was supported by the National Key Research and Development Program of China grant no. 2021YFF1200902 (R.J.), the National Natural Science Foundation of China grants nos. 61873141 (R.J.), 61721003 (X.Z.), 62273194 (R.J.), 62203236 (S.C.).

## Author contributions

R.J. and S.C conceived the study and supervised the project. Z.L. and S.C. designed, implemented and validated PAST. X.C. and X.Z. helped with analysing the results. S.C., Z.L.

X.Z. and R.J. wrote the manuscript, with input from all the authors.

