## supplementary materials for "PAST: latent feature extraction with a Prior-based self-Attention framework for Spatial Transcriptomics"

**Algorithm 1** Construction of pseudo-bulk gene expression matrix

**Input:** Preprocessed reference gene expression matrix $\boldsymbol{R}$, spatial domain annotation vector $\boldsymbol{Y}$, minimum pseudo-bulk samples $N_m$, sampling ratio $r$

**Output:** Pseudo-bulk gene expression matrix $\widetilde{\boldsymbol{R}}$

**1:** Initialize an empty pseudo-bulk matrix $\widetilde{\boldsymbol{R}} = \{\}$

**2: for** $c = 1,2, \dots, K$ **do** /* $K$ is the number of categories in the spatial domain annotation vector $\boldsymbol{Y}$ */

**3:** Obtain $\Omega_c = \{\boldsymbol{s} | \boldsymbol{s} \in \boldsymbol{R}, Cat(\boldsymbol{s}) = \boldsymbol{Y}_c\}$ /*$Cat(\boldsymbol{s})$ denotes the spatial domain of spot $\boldsymbol{s}$, $\boldsymbol{Y}_c$ denotes the $c$-th category in spatial domain annotation, $\Omega_c$ denotes the set of spots belonging to the $c$-th category */

**4:** Calculate pseudo-bulk sample $\boldsymbol{S}_c = Avg(\Omega_c)$, then Add $\boldsymbol{S}_c$ to $\widetilde{\boldsymbol{R}}$ /* $Avg(\Omega_c)$ denotes the average expression of all spots in $\Omega_c$ */

**5: end for**

**6: while** $|\widetilde{\boldsymbol{R}}| < N_m$ **do** /* $|\widetilde{\boldsymbol{R}}|$ denotes the number of samples in $\widetilde{\boldsymbol{R}}$ */

**7: for** $c = 1,2, \dots, K$ **do**

**8:** Randomly select spots from $\Omega_c = \{\boldsymbol{s} | \boldsymbol{s} \in \boldsymbol{R}, Cat(\boldsymbol{s}) = \boldsymbol{Y}_c\}$ with sampling ratio $r$ to obtain $\Omega_{c,r}$

**9:** Calculate pseudo-bulk sample $\boldsymbol{S}_{c,r} = Avg(\Omega_{c,r})$, then add $\boldsymbol{S}_{c,r}$ to $\widetilde{\boldsymbol{R}}$

**10: end for**

**11: end while**

**12: return** $\widetilde{\boldsymbol{R}}$

**Algorithm 2** Ripple walk sampler

**Input:** Graph $\mathcal{G} = (\mathcal{V}, \mathcal{E})$, expansion ratio $r$, batch size $B$ /* $\mathcal{V}$ denotes the set of spots in ST datasets, while $\mathcal{E}$ denotes the set of edges between spots */

**Output:** Subgraph $\mathcal{G}_k$

**1:** Initialize $\mathcal{G}_k = (\mathcal{V}_k, \mathcal{E}_k)$ where $\mathcal{V}_\mathrm{k} = \emptyset$

**2:** Randomly select the initial node $v_s$, then add $v_s$ into $\mathcal{G}_k$

**3: while** $|\mathcal{V}_k| < B$ **do**

**4:** Obtain $Neigh(\mathcal{V}_k) = \{n|(n,j) \in \mathcal{E}, j \in \mathcal{V}_k, n \in \mathcal{V}\backslash\mathcal{V}_k\}$ /* obtain neighborhood set of $\mathcal{V}_k$ */

**5:** Randomly select spots from $Neigh(\mathcal{V}_k)$ with ratio $r$, then add them into $\mathcal{V}_k$

**6: end while**

**7: return** $\mathcal{G}_k$

**Algorithm 3** Ripple walk training

**Input:** Untrained PAST model $H_{\boldsymbol{W}}(\cdot)$, target gene expression matrix $\boldsymbol{M}$, $k$-NN graph $\boldsymbol{\mathcal{G}}$, expansion ratio $r$, batch size $B$, subgraph number within one training iteration $S$, training iteration number $T$

**Output:** Trained PAST model $\widetilde{H}_{\boldsymbol{W}}(\cdot)$

**1:** Initialize mini-batch subgraph set $U = \{\}$

**2: for** $k = 1,2,\dots,S$ **do**

**3:** Obtain subgraph $\boldsymbol{\mathcal{G}}_k$ using ripple walk sampler with expansion ratio $r$ and batch size $B$ /* **Algorithm 2** */

**4:** Add $\boldsymbol{\mathcal{G}}_k$ to $U$

**5: end for**

**6: for** $i = 1,2,\dots,T$ **do**

**7: for** $k = 1,2,\dots,S$ **do**

**8:** Select subgraph $\boldsymbol{\mathcal{G}}_k$ from $U$ and the corresponding gene expression matrix $\boldsymbol{M}_k$ from $\boldsymbol{M}$ /* $\boldsymbol{M}_k$ is the corresponding subset of gene expression matrix $\boldsymbol{M}$ */

**9:** PAST model $H_{\boldsymbol{W}}(\cdot)$ performs forward and back propagation based on the subgraph $\boldsymbol{\mathcal{G}}_k$ and corresponding gene expression matrix $\boldsymbol{M}_k$

**10:** Update the parameters $\boldsymbol{W}$ of PAST model according to the gradient of the loss function

**11: end for**

**12: end for**

**13: return** $\widetilde{H}_{\boldsymbol{W}}(\cdot)$

**Algorithm 4** Ripple walk prediction

**Input:** Trained PAST model $\widetilde{H}_{\boldsymbol{W}}(\cdot)$, gene expression matrix $\boldsymbol{M}$, k-NN graph $\boldsymbol{\mathcal{G}} = (\mathcal{V}, \mathcal{E})$, expansion ratio $r$, batch size $B$

**Output:** Latent embedding matrix $\widetilde{\boldsymbol{M}}$

**1:** Initialize mini-batch subgraph set $U = \{\}$, sampled node set $L = \{\}$

**2: while** $\mathcal{V} \backslash L \neq \emptyset$ **do**

**3:** Obtain subgraph $\boldsymbol{\mathcal{G}}_k = (\mathcal{V}_k, \mathcal{E}_k)$ using Refined ripple walk sampler with expansion ratio $r$ and batch size $B$ /* Refined ripple walk sampler randomly select initial nodes from non-sampled node set $\mathcal{V} \backslash L$ instead of $\mathcal{V}$ in step 2 of **Algorithm 2** */

**4:** Update sampled node set $L = L \cup \mathcal{V}_k$

**5:** Add $\boldsymbol{\mathcal{G}}_k$ to $U$

**6: end while**

**7: for** $k = 1,2, \dots, |U|$ **do**

**8:** Select subgraph $\boldsymbol{\mathcal{G}}_k$ from $U$ and the corresponding gene expression matrix $\boldsymbol{M}_k$ from $\boldsymbol{M}$

**9:** Obtain latent embeddings $\widetilde{\boldsymbol{M}}_k$ with trained PAST model $\widetilde{H}_{\boldsymbol{W}}(\cdot)$ based on the subgraph $\boldsymbol{\mathcal{G}}_k$ and corresponding gene expression matrix $\boldsymbol{M}_k$

**10: end for**

**11:** Concatenate all the $\widetilde{\boldsymbol{M}}_k$ to obtain $\widetilde{\boldsymbol{M}}_{\text{concat}}$, and then calculate the average latent embeddings for samples contained in multiple subgraph in $\widetilde{\boldsymbol{M}}_{\text{concat}}$ to get latent embedding matrix $\widetilde{\boldsymbol{M}}$

**13: return** $\widetilde{\boldsymbol{M}}$

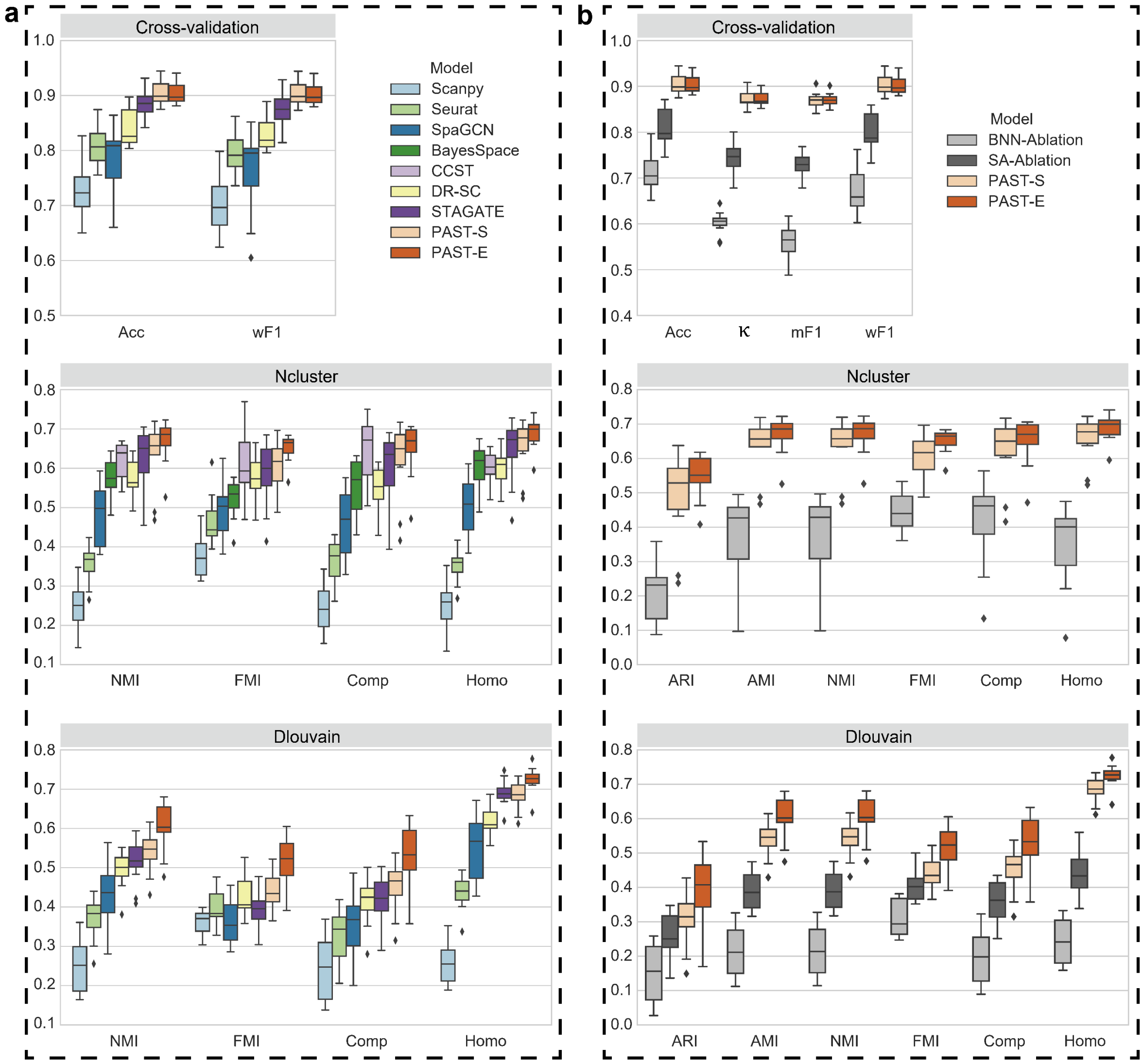

**Fig. S1 | Quantitative performance evaluation of spatial domain characterization on the 10x Visium dorsolateral prefrontal cortex (DLPFC) datasets.** Quantitative evaluation via supervised cross-validation and unsupervised spatial clustering with a specified number of clusters (Ncluster) and with default resolution (Dlouvain). The center line, box limits and whiskers in the boxplots are the median, upper and lower quartiles, and 1.5× interquartile range, respectively. **a**, Evaluation of PAST and other baseline methods. The cross-validation performance was evaluated by the average score of accuracy (Acc) and weighted F1 score (wF1), respectively, in the 5-fold experiments. The spatial clustering performance was evaluated by normalized mutual information (NMI), fowlkes-mallows index (FMI), completeness (Comp) and homogeneity (Homo). **b**, Evaluation of PAST and ablation models. We replace the BNN module (BNN-Ablation) and self-attention mechanism (SA-Ablation) in PAST model with fully connected layer for ablation study. Note that SA-Ablation model encountered NULL-TYPE error when performing mclust algorithm with a specified number of clusters. The cross-validation performance was evaluated by the average score of accuracy (Acc), Cohen's kappa value (κ), mean F1 score (mF1) and weighted F1 score (wF1), respectively, in the 5-fold experiments. The spatial clustering performance was evaluated by adjusted rand index (ARI), adjusted mutual information (AMI), normalized mutual information (NMI), fowlkes-mallows index (FMI), completeness (Comp) and homogeneity (Homo).

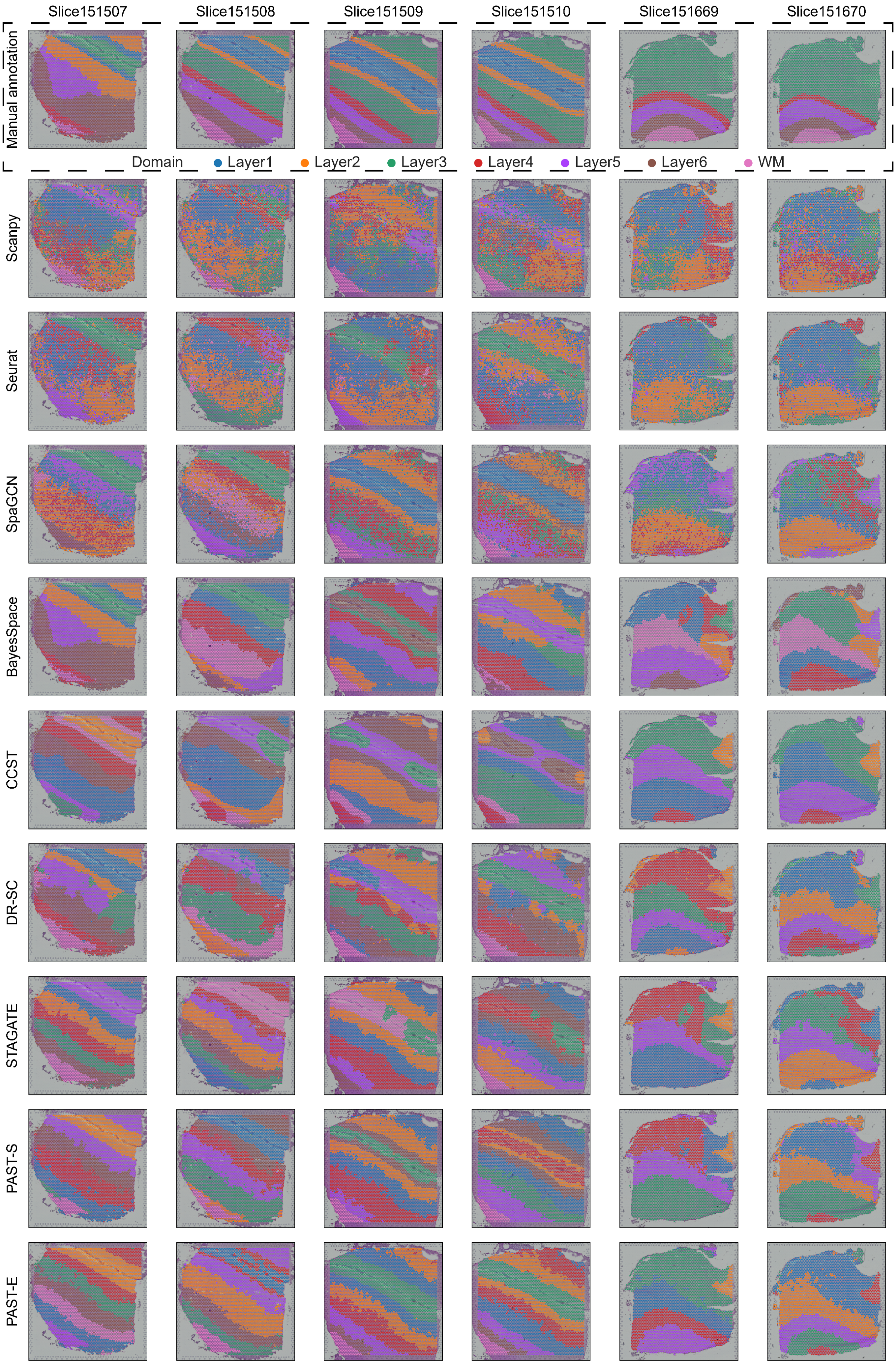

Pick up on the next page

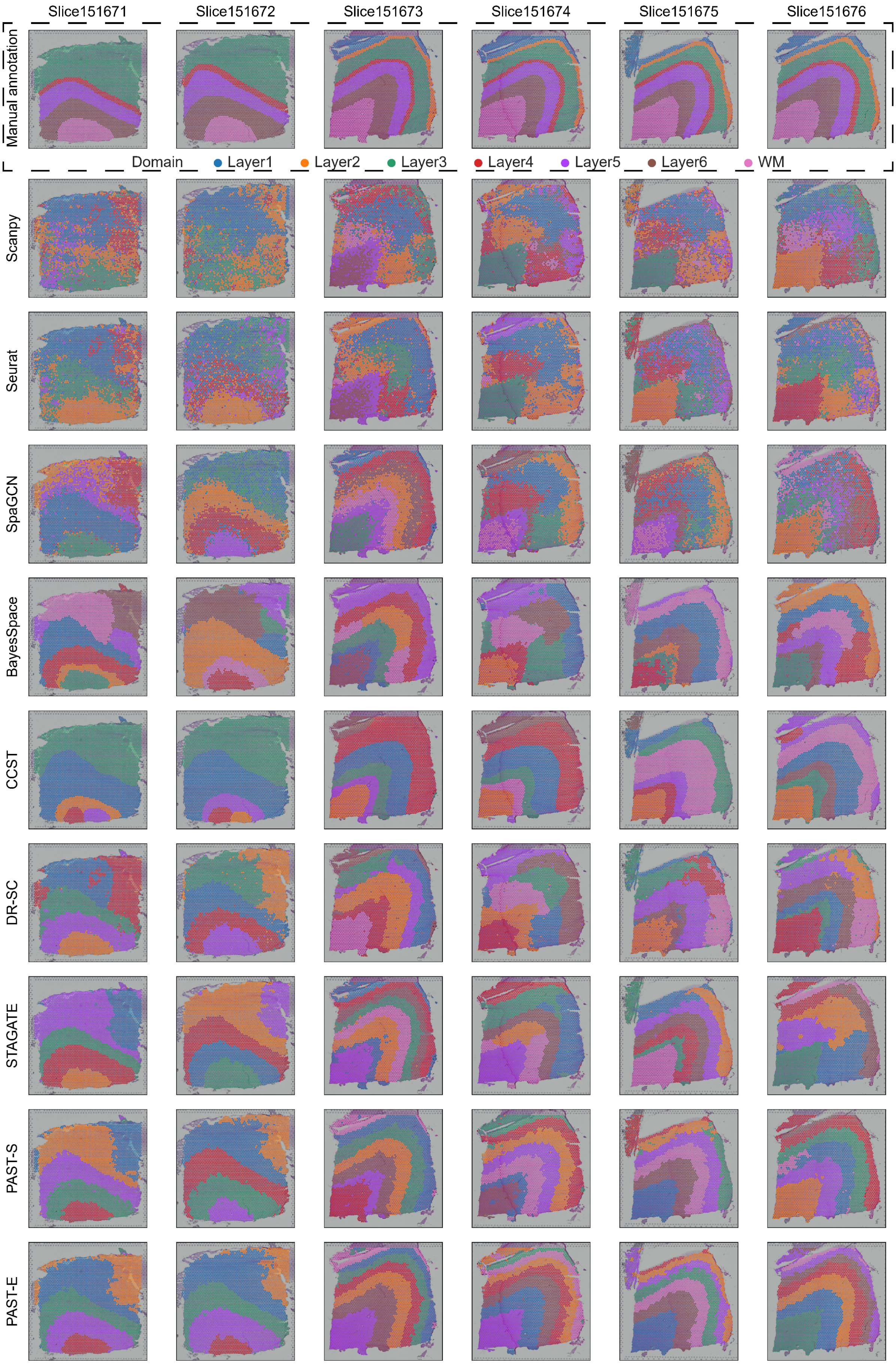

**Fig. S2 | Spatial clustering results on 10x Visium DLPFC datasets.**

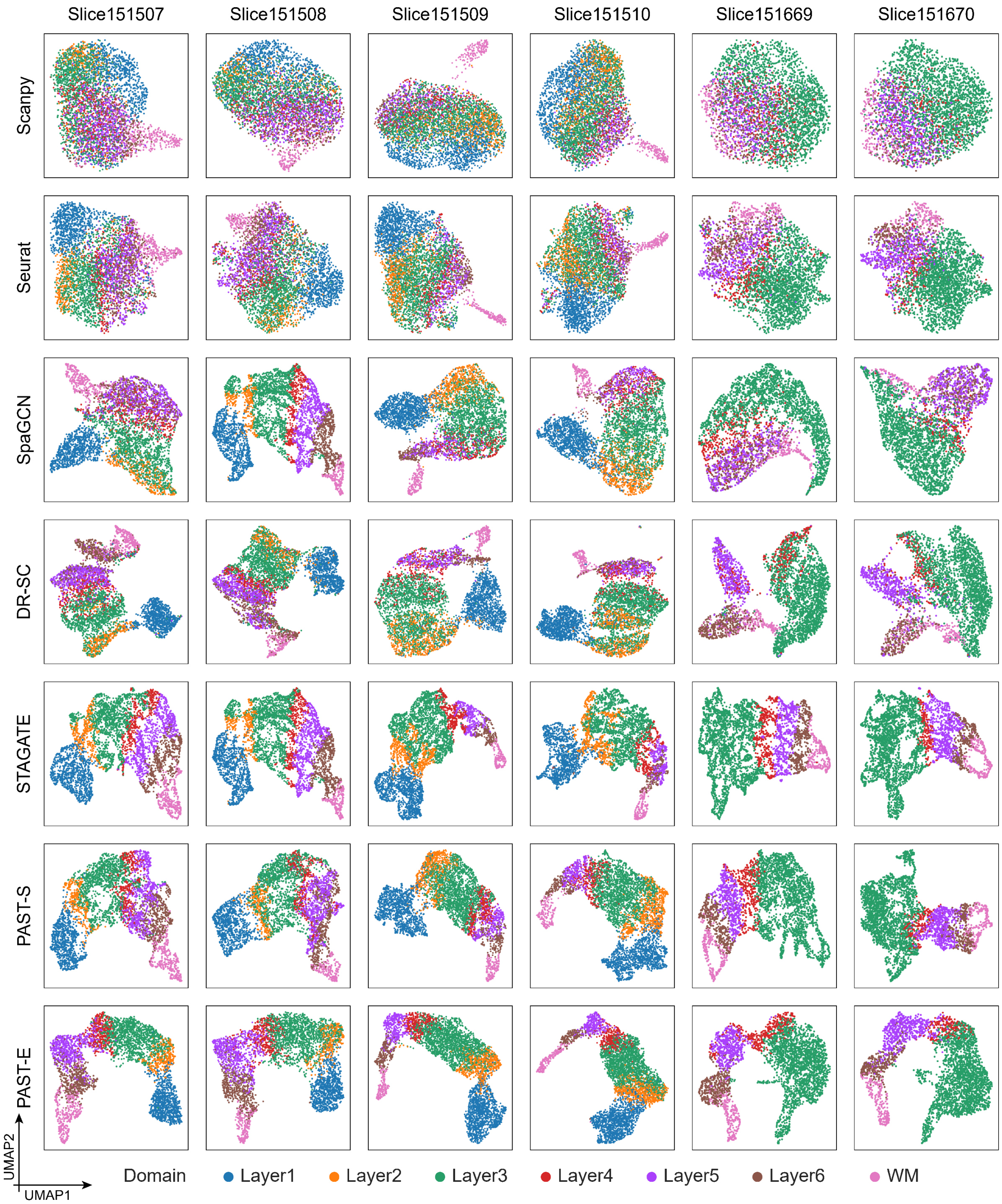

Pick up on the next page

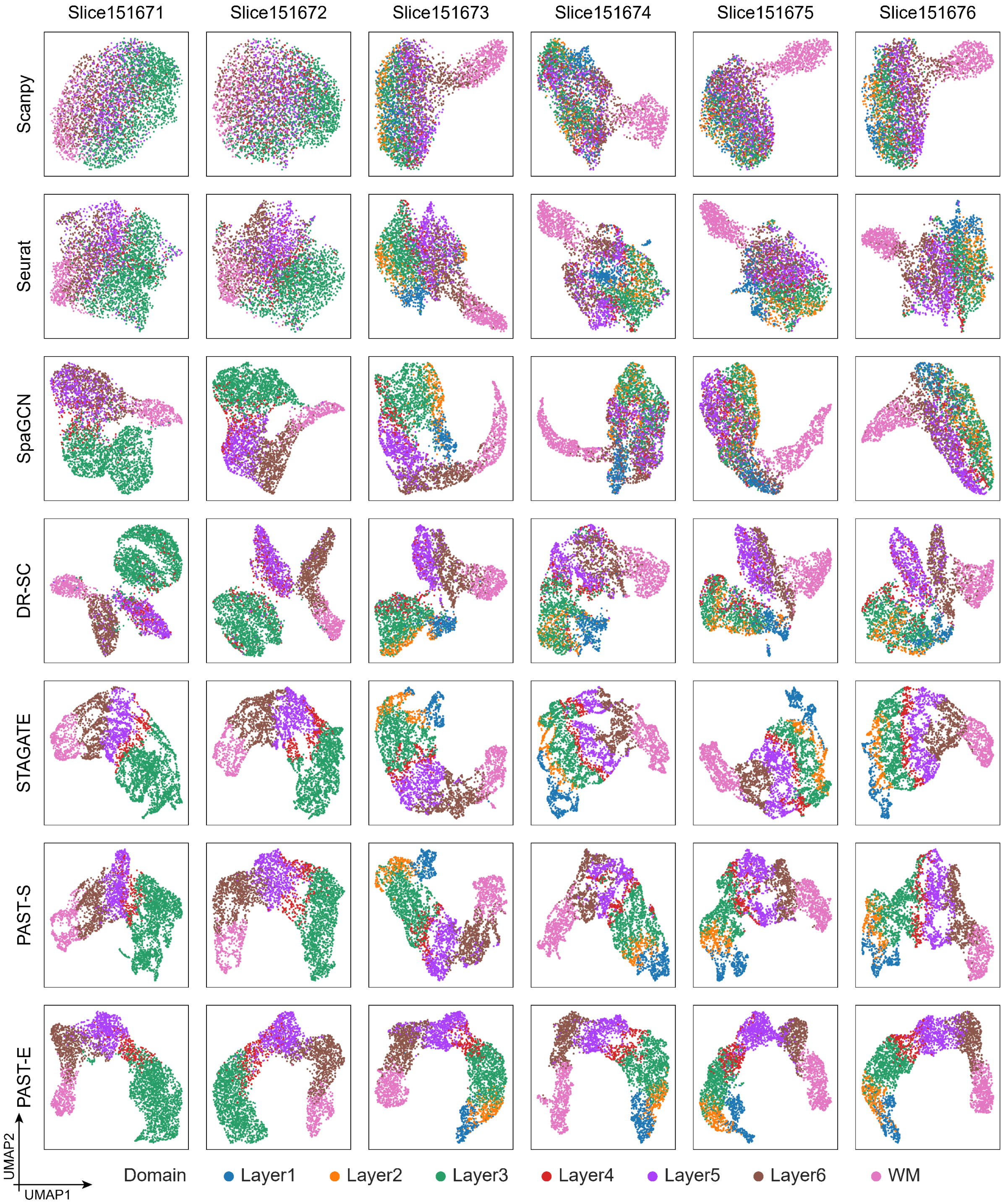

**Fig. S3 | UMAP visualization on 10x Visium DLPFC datasets.**

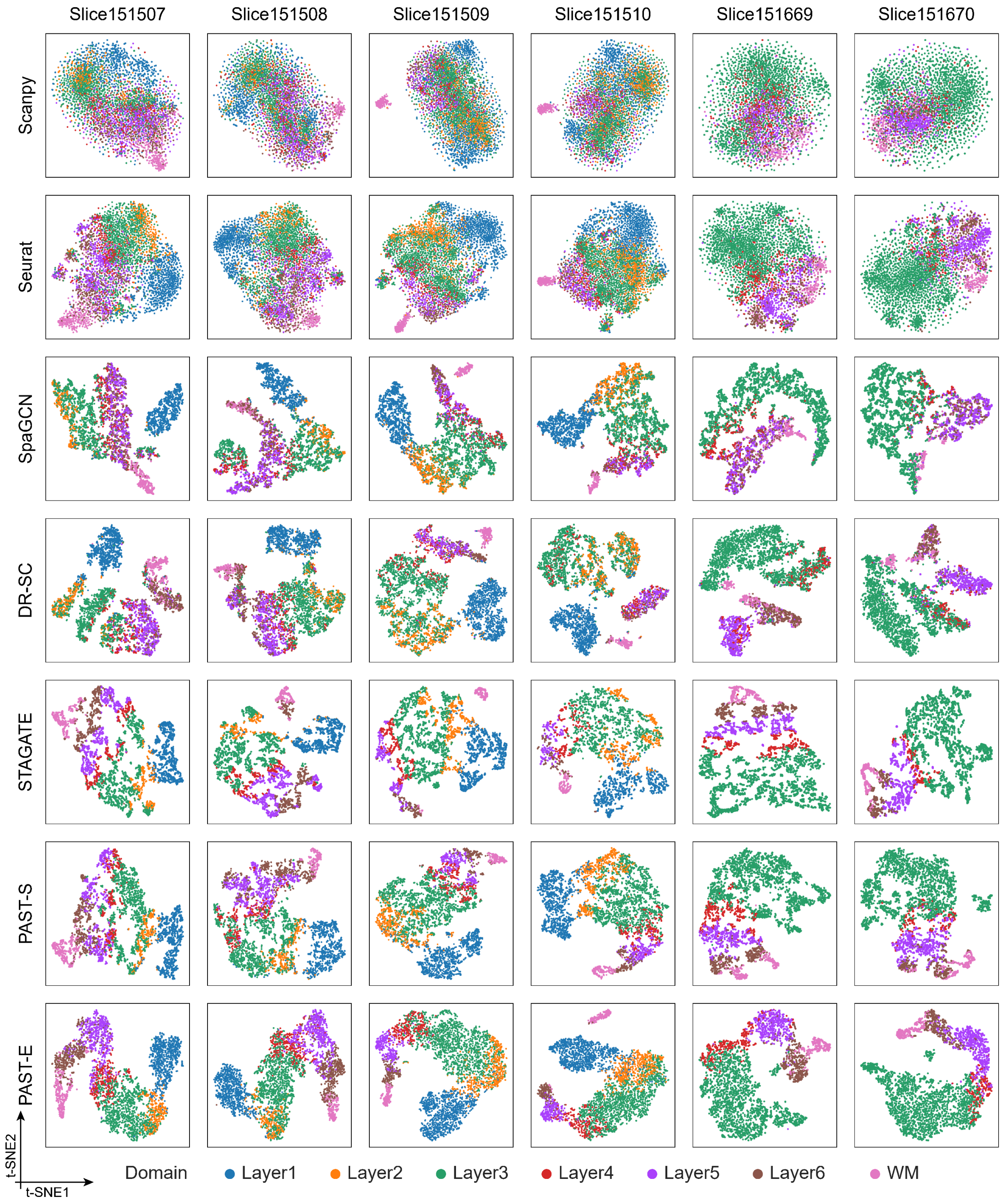

Pick up on the next page

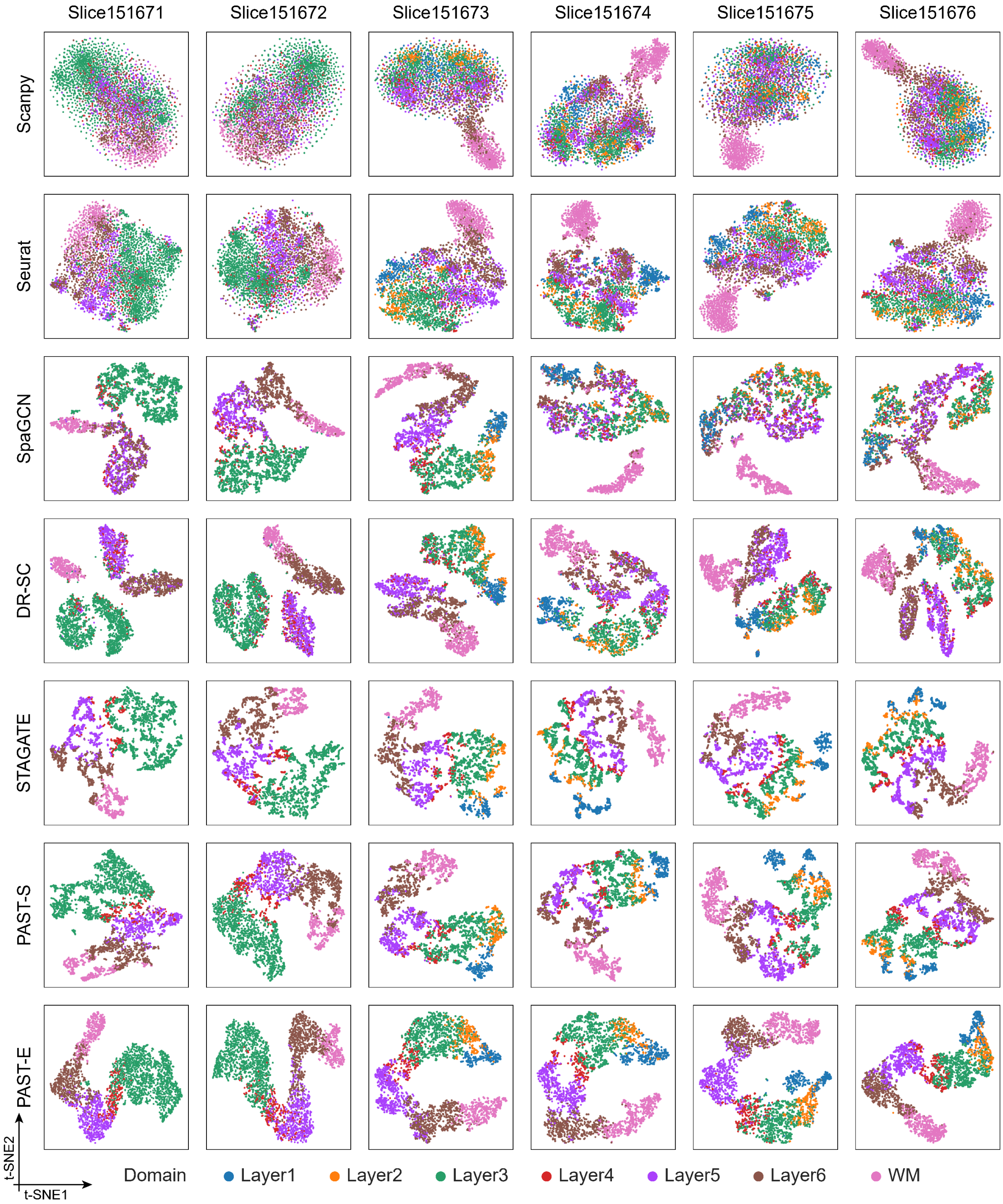

**Fig. S4 | t-SNE visualization on 10x Visium DLPFC datasets.**

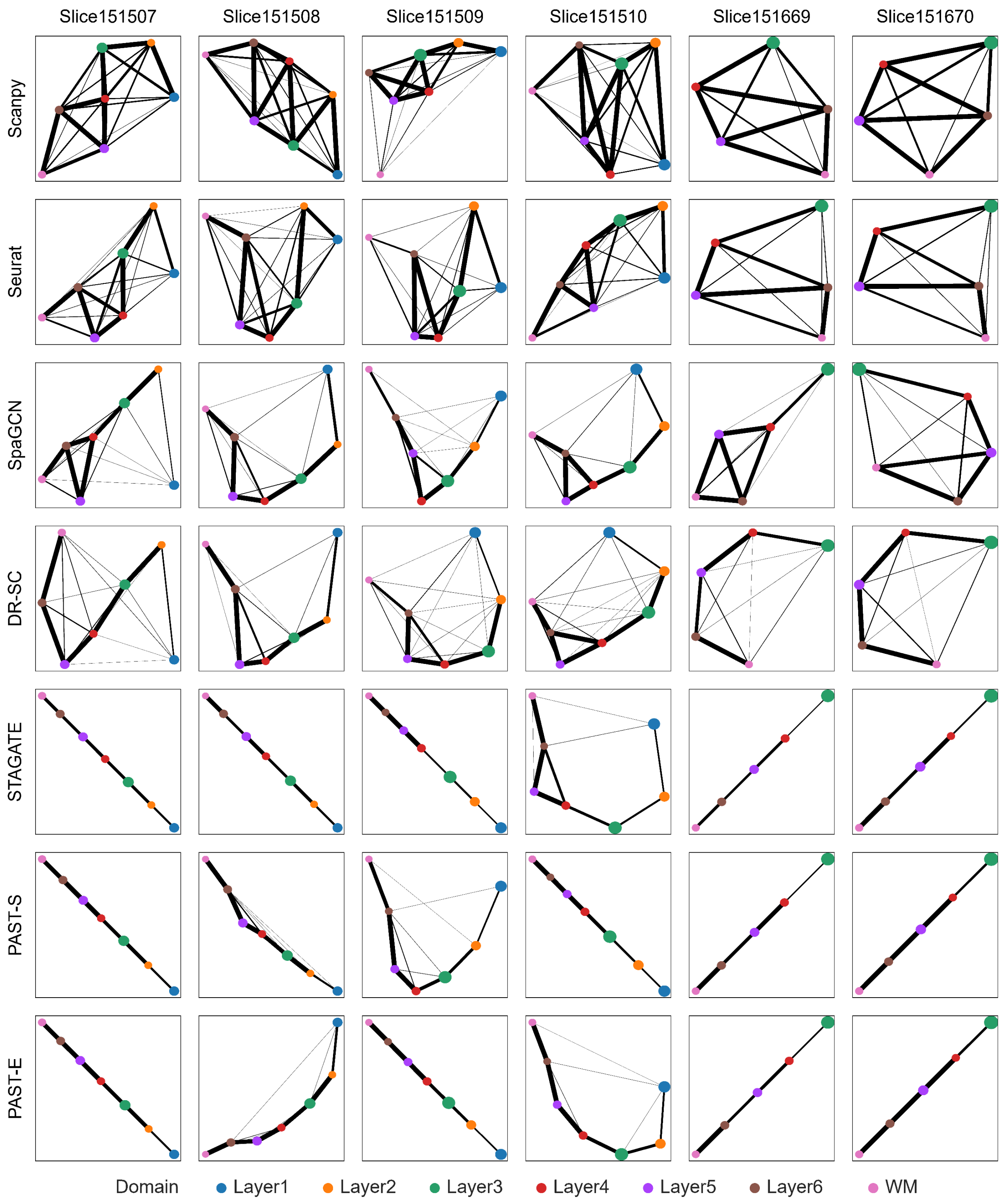

Pick up on the next page

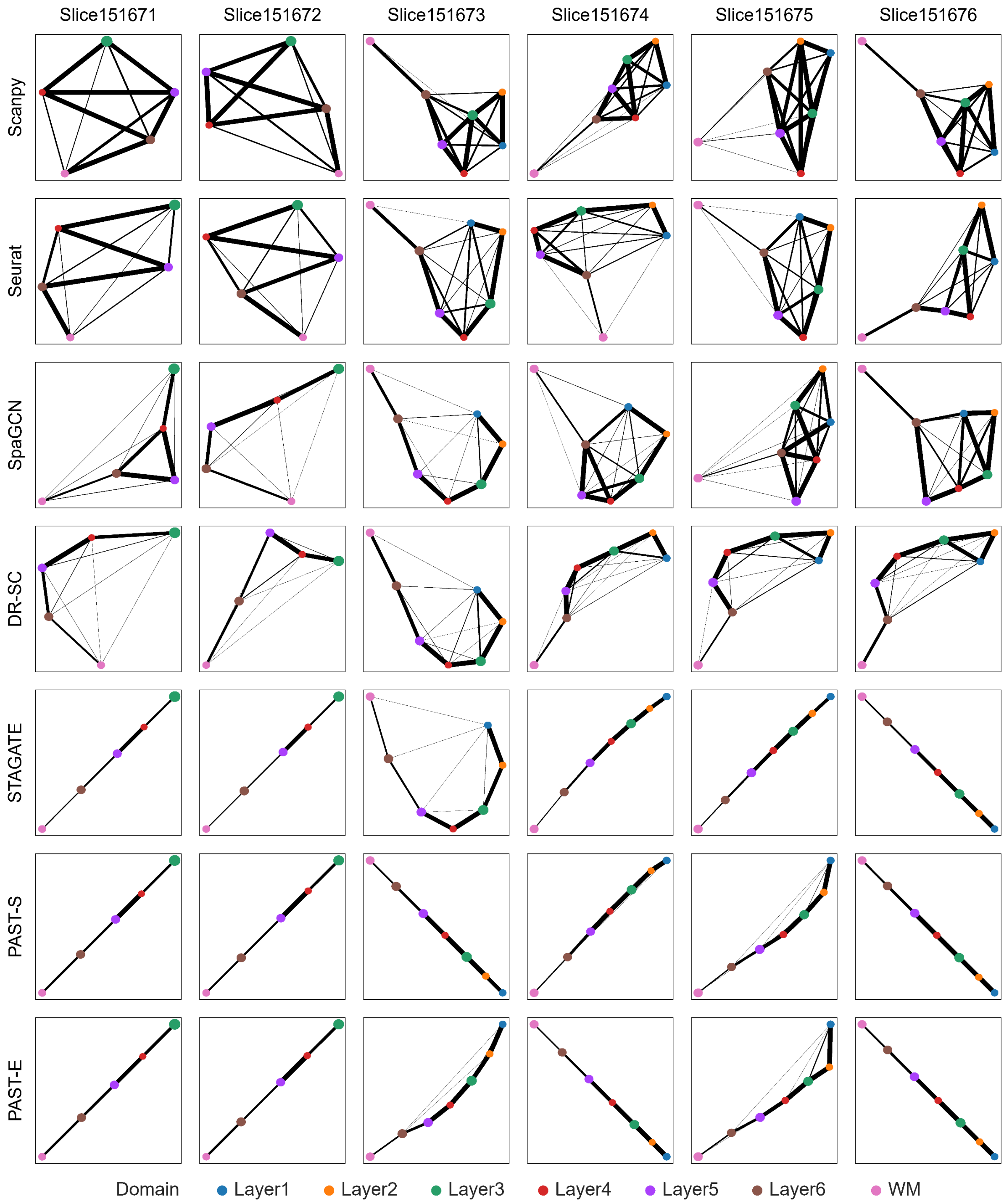

**Fig. S5 | PAGA trajectory inference results on 10x Visium DLPFC datasets.**

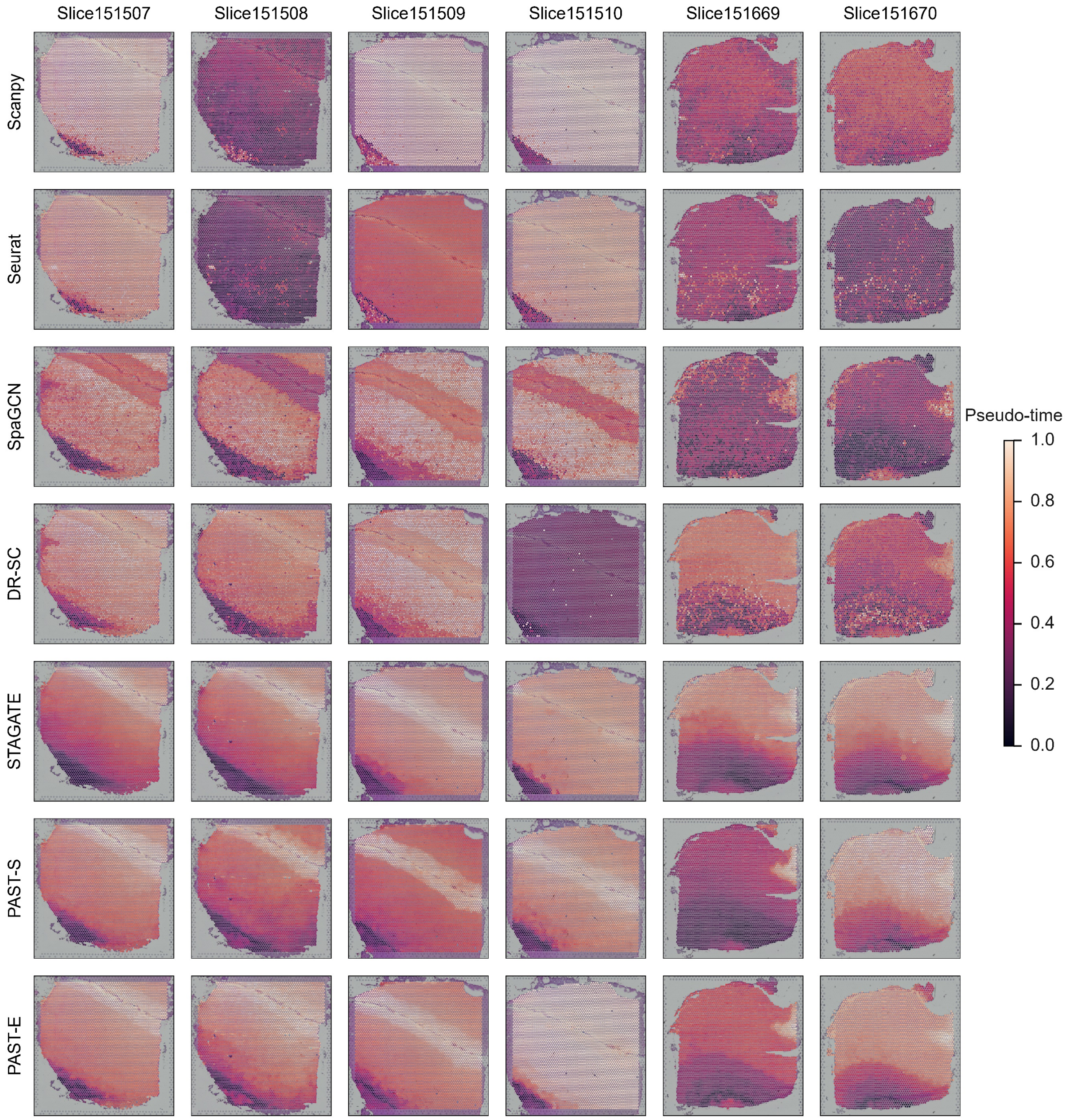

Pick up on the next page

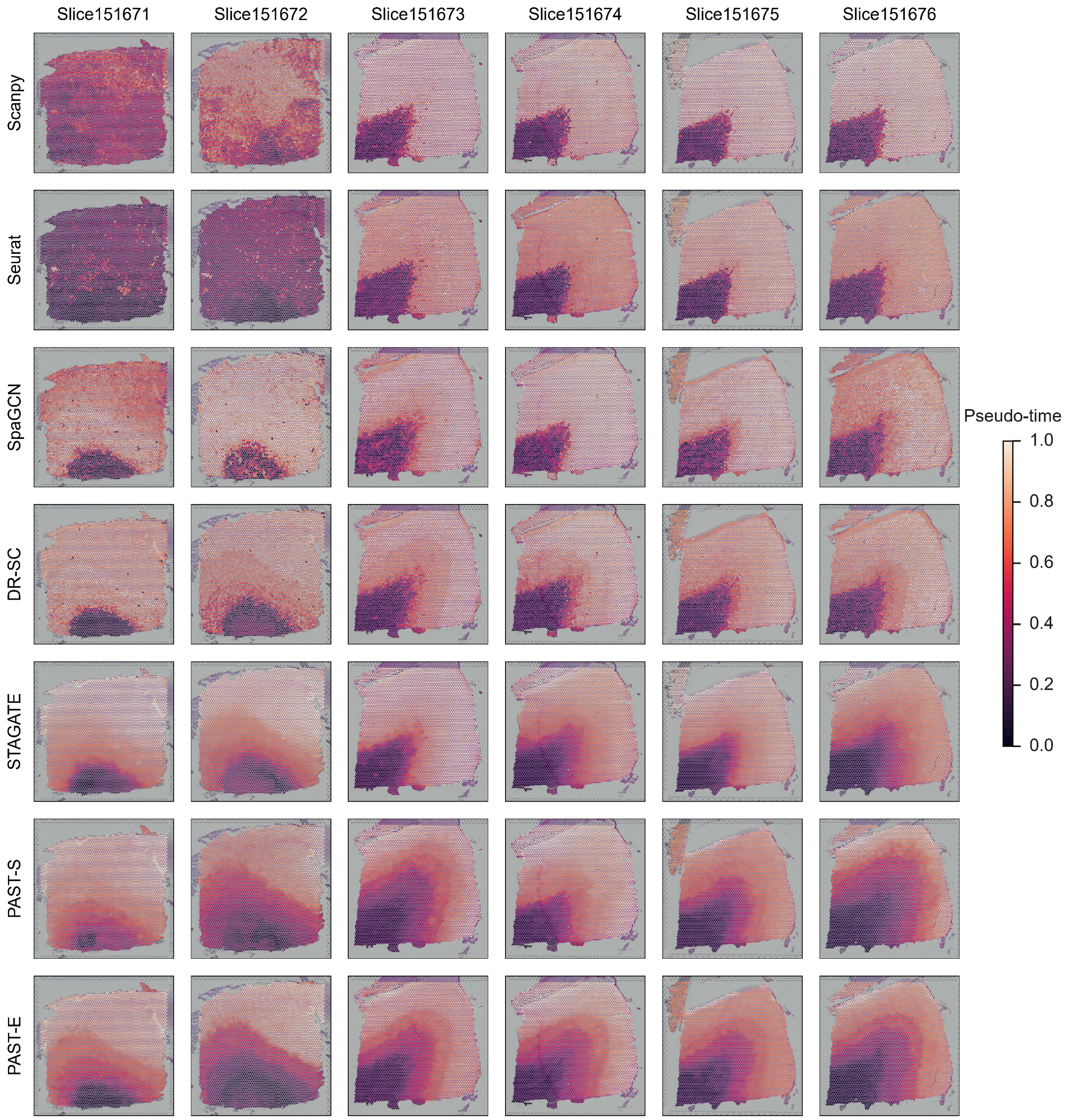

**Fig. S6 | DPT pseudo-time analysis on 10x Visium DLPFC datasets.**

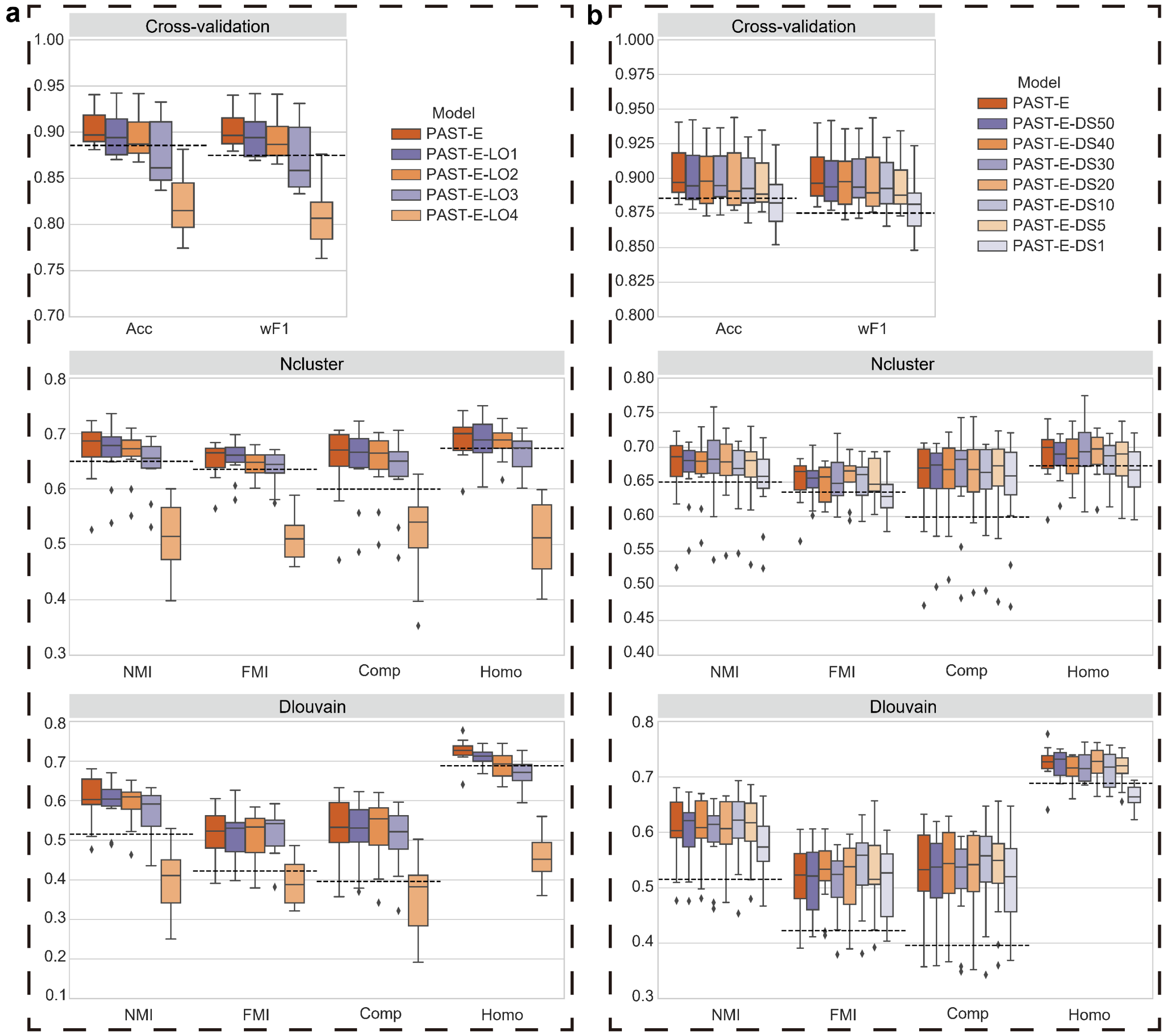

**Fig. S7 | The performance of PAST with incomplete or different scales of external reference on 10x Visium DLPFC datasets.** Quantitative performance evaluation of spatial domain characterization on 12 DLPFC slices via supervised cross-validation and unsupervised spatial clustering with a specified number of clusters (Ncluster) and with default resolution (Dlouvain). The cross-validation performance was evaluated by the average score of Acc and wF1, respectively, in the 5-fold experiments. The spatial clustering performance was evaluated by NMI, FMI, Comp and Homo. The horizontal dashed lines denote the corresponding median scores of the second-best method, STAGATE. The center line, box limits and whiskers in the boxplots are the median, upper and lower quartiles, and 1.5× interquartile range, respectively. **a,** The spatial domains in external reference data of PAST were gradually left out. **b,** The spots in external reference data of PAST were downsampled to construct reference data with various scales.

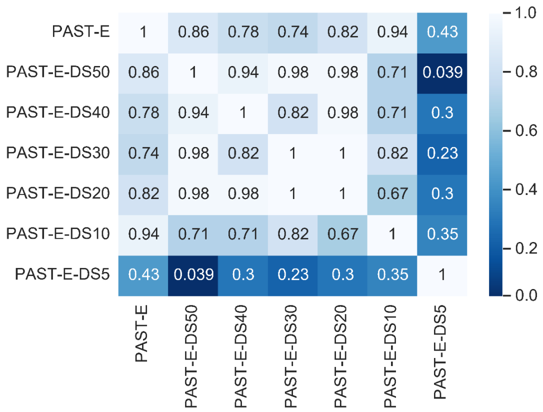

**Fig. S8 | Two-sided Wilcoxon signed-rank tests on PAST with different scales of external reference on 10x Visium DLPFC datasets.** We utilized 16 metrics to evaluate PAST with different scales of external reference, i.e., 4 metrics for cross validation, 6 metrics for Ncluster and 6 metrics for Dlouvain. Specifically, we calculated median scores on 12 DLPFC slices, and conducted two-sided Wilcoxon signed-rank tests based on the 16 metrics to explore the impact of different scales of external reference on the performance of PAST.

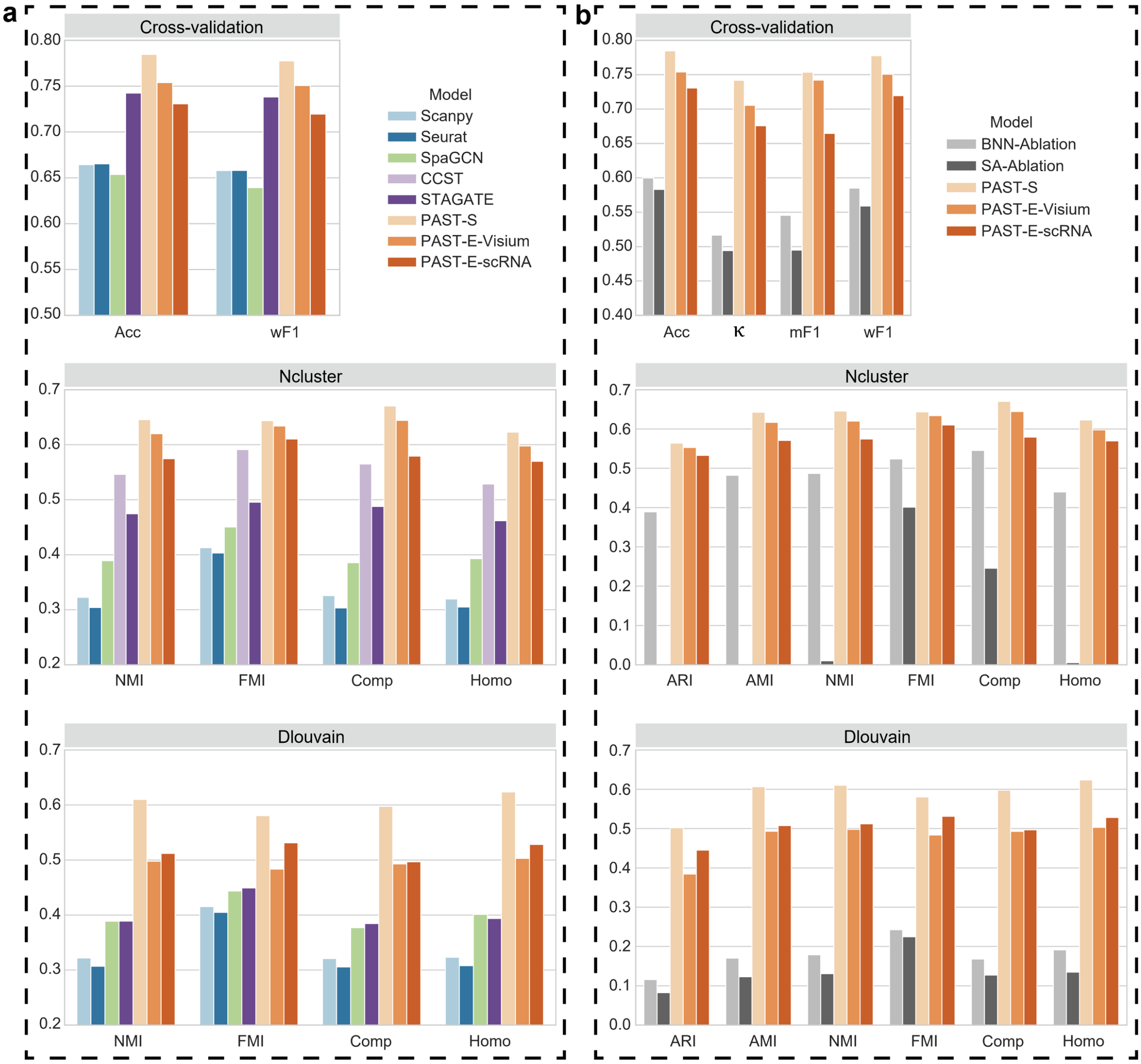

**Fig. S9 | Quantitative performance evaluation of spatial domain characterization on the STARmap mouse primary visual cortex (MPVC) dataset.** Quantitative evaluation via supervised cross-validation and unsupervised spatial clustering with a specified number of clusters (Ncluster) and with default resolution (Dlouvain). **a**, Evaluation of PAST and other baseline methods. The cross-validation performance was evaluated by the average score of Acc and wF1, respectively, in the 5-fold experiments. The spatial clustering performance was evaluated by NMI, FMI, Comp and Homo. **b**, Evaluation of PAST and ablation models. The cross-validation performance was evaluated by the average score of Acc, κ, mF1 and wF1, respectively, in the 5-fold experiments. The spatial clustering performance was evaluated by ARI, AMI, NMI, FMI, Comp and Homo.

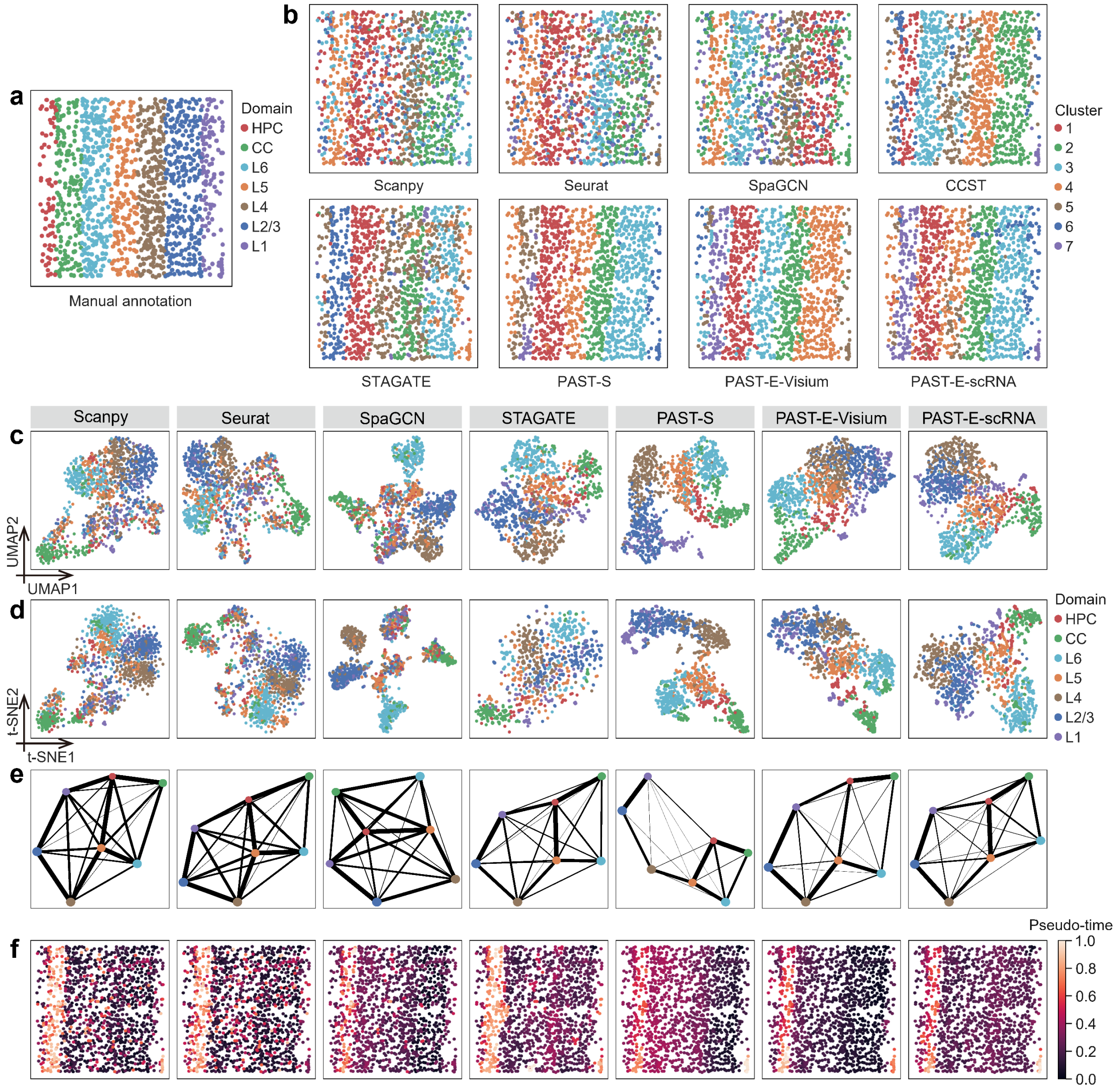

**Fig. S10 | Spatial clustering, UMAP visualization, t-SNE visualization, PAGA and DPT pseudo-time analysis results on STARmap MPVC dataset. a**, Manual annotation of MPVC. **b**, Spatial clustering results **c**, UMAP visualization **d**, t-SNE visualization **e**, PAGA trajectory inference results and **f**, DPT pseudo-time analysis results on MPVC.

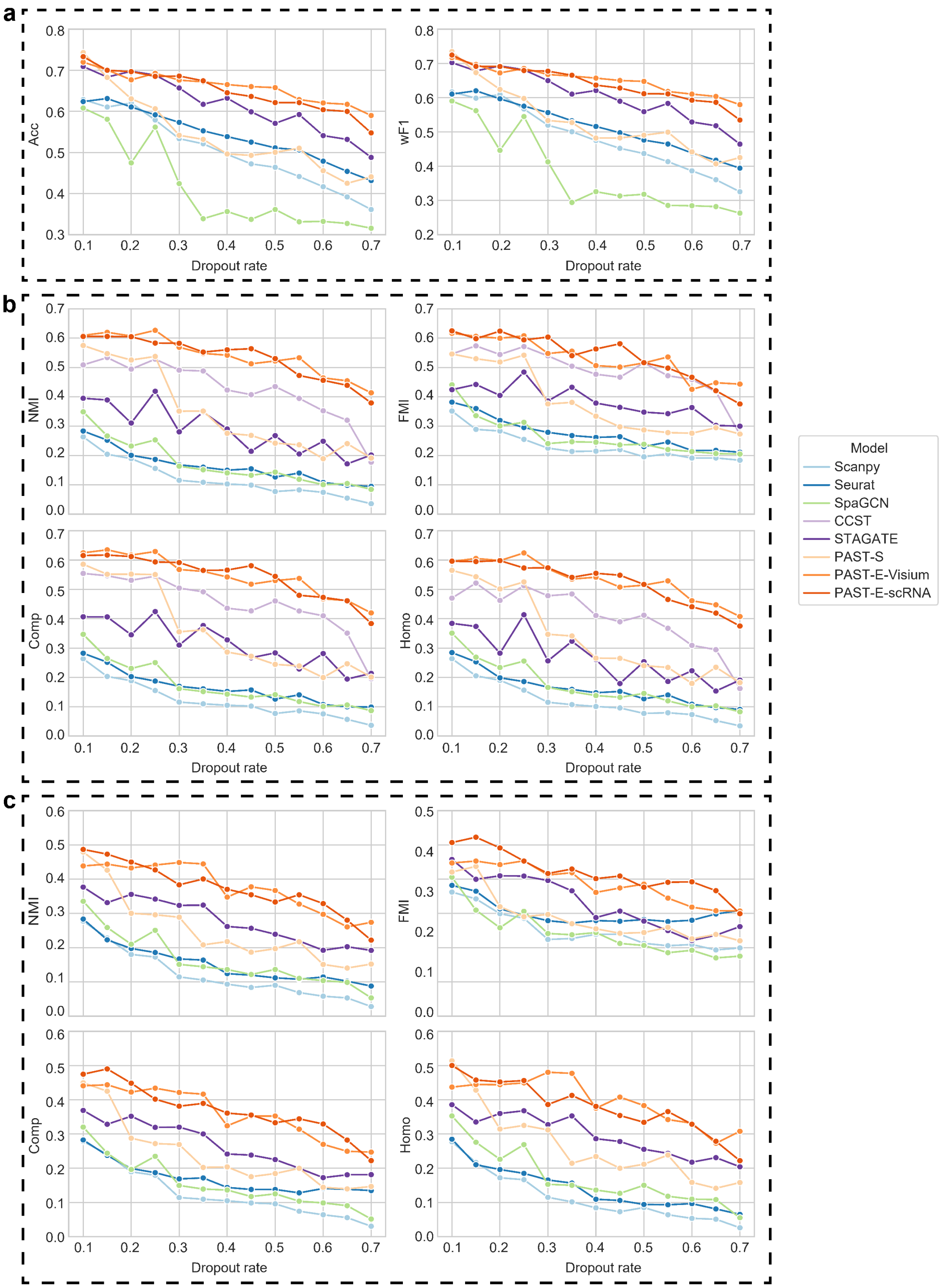

**Fig. S11 | The performance of different methods under various dropout rates on STARmap MPVC dataset.** We added different levels of dropout noise into target gene expression data and evaluated the performance of different methods in terms of **a**, cross-validation on latent embeddings evaluated by Acc and wF1 and spatial clustering with **b**, a specified number of clusters (Ncluster) or **c**, default resolution (Dlouvain) evaluated by NMI, FMI, Comp and Homo.

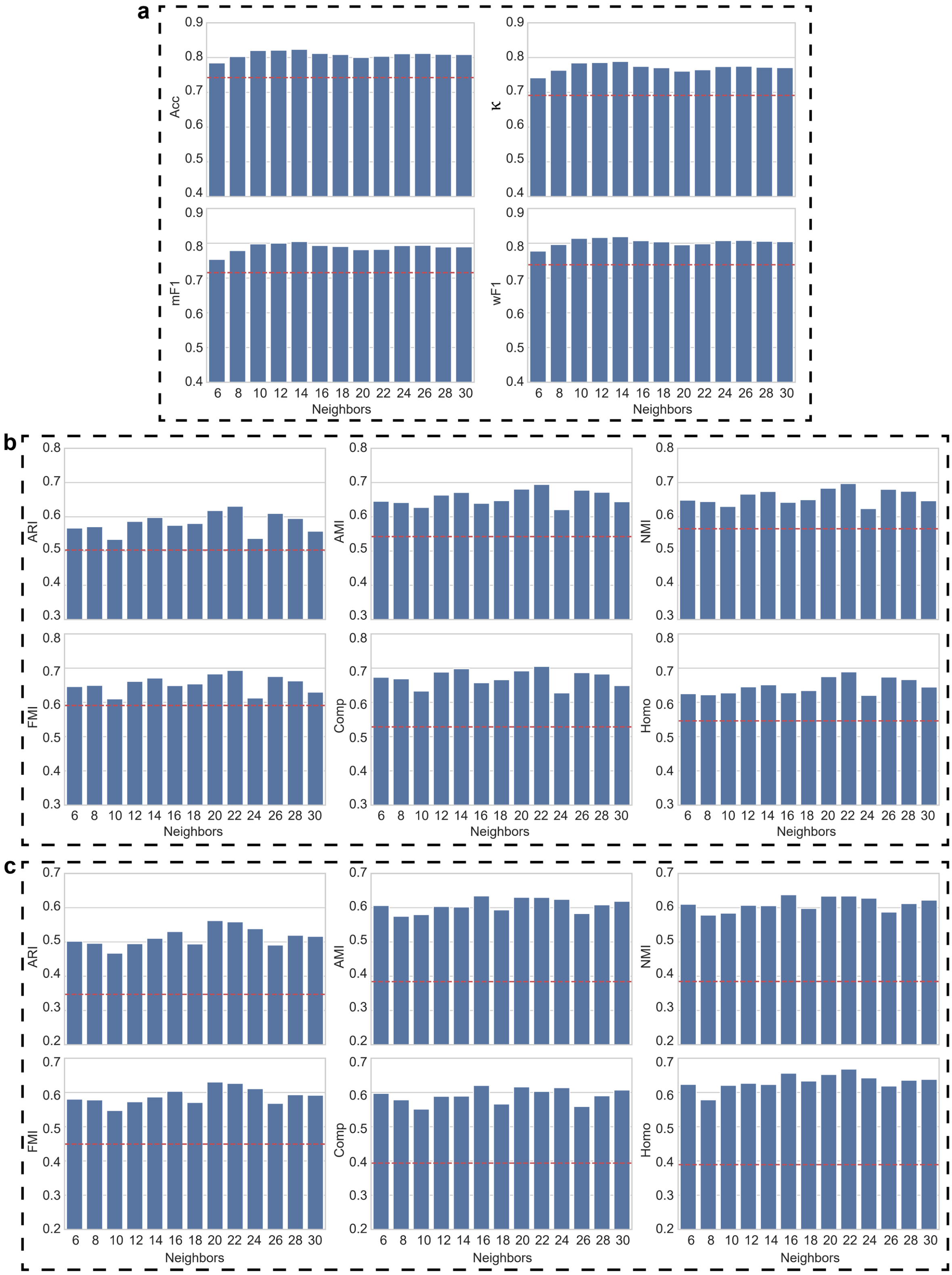

**Fig. S12 | The performance of PAST with different neighbors on STARmap MPVC dataset.** We changed the number of neighbors in self-attention mechanism to explore the robustness of PAST. **a**, Supervised cross-validation evaluated by Acc, κ, mF1 and wF1. The red lines denote the performance of the second-best method, STAGATE. **b**, Spatial clustering with a specified number of clusters (Ncluster) evaluated by ARI, AMI, NMI, FMI, Comp and Homo. The red lines denote the performance of the second-best method, CCST. **c**, Spatial clustering with default resolution (Dlouvain) evaluated by ARI, AMI, NMI, FMI, Comp and Homo. The red lines denote the performance of the second-best method, STAGATE.

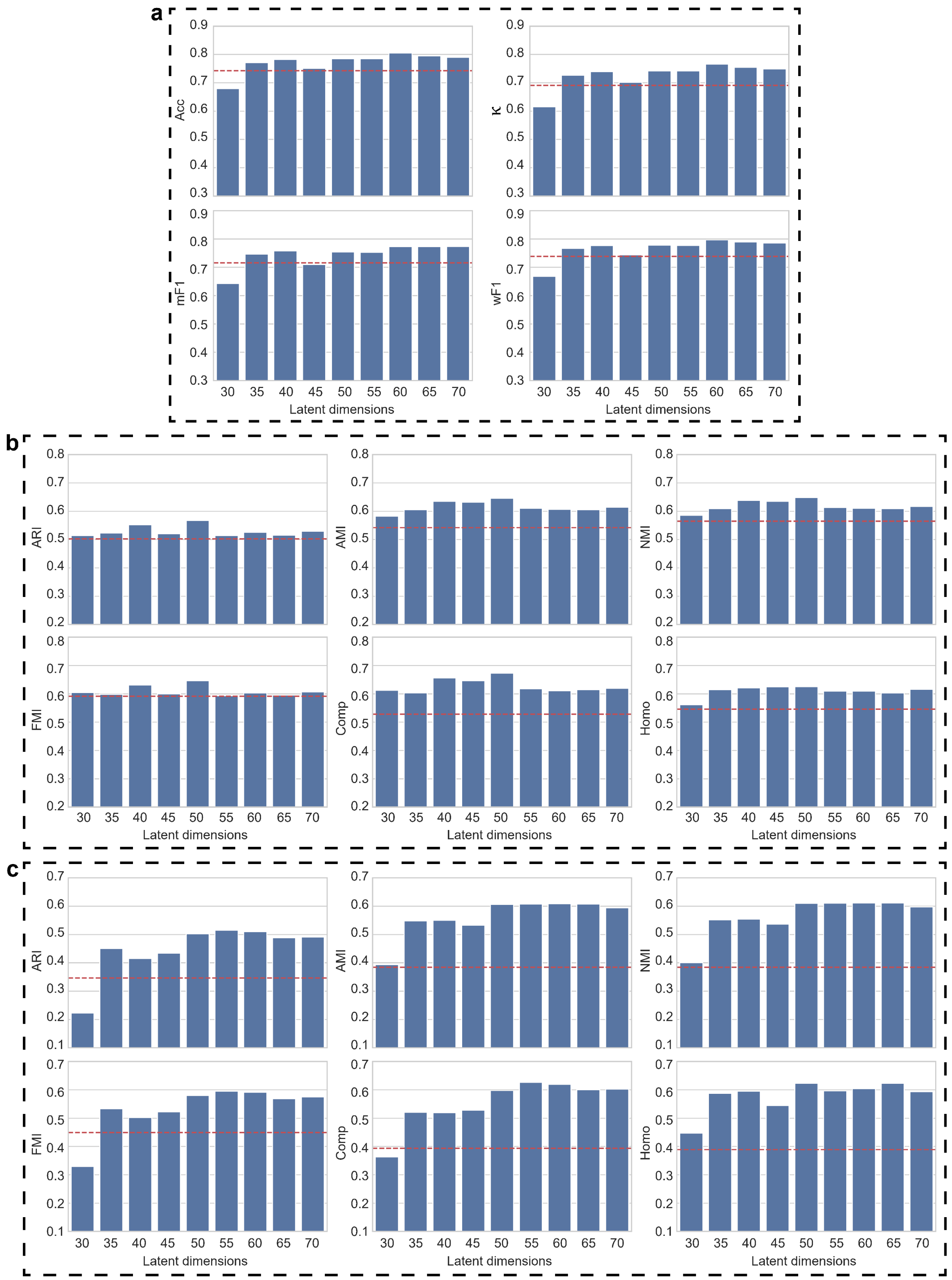

**Fig. S13 | The performance of PAST with different latent dimensions on STARmap MPVC dataset.** We changed the number of latent dimensions to explore the robustness of PAST. **a**, Supervised cross-validation evaluated by Acc, κ, mF1 and wF1. The red lines denote the performance of the second-best method, STAGATE. **b**, Spatial clustering with a specified number of clusters (Ncluster) evaluated by ARI, AMI, NMI, FMI, Comp and Homo. The red lines denote the performance of the second-best method, CCST. **c**, Spatial clustering with default resolution (Dlouvain) evaluated by ARI, AMI, NMI, FMI, Comp and Homo. The red lines denote the performance of the second-best method, STAGATE.

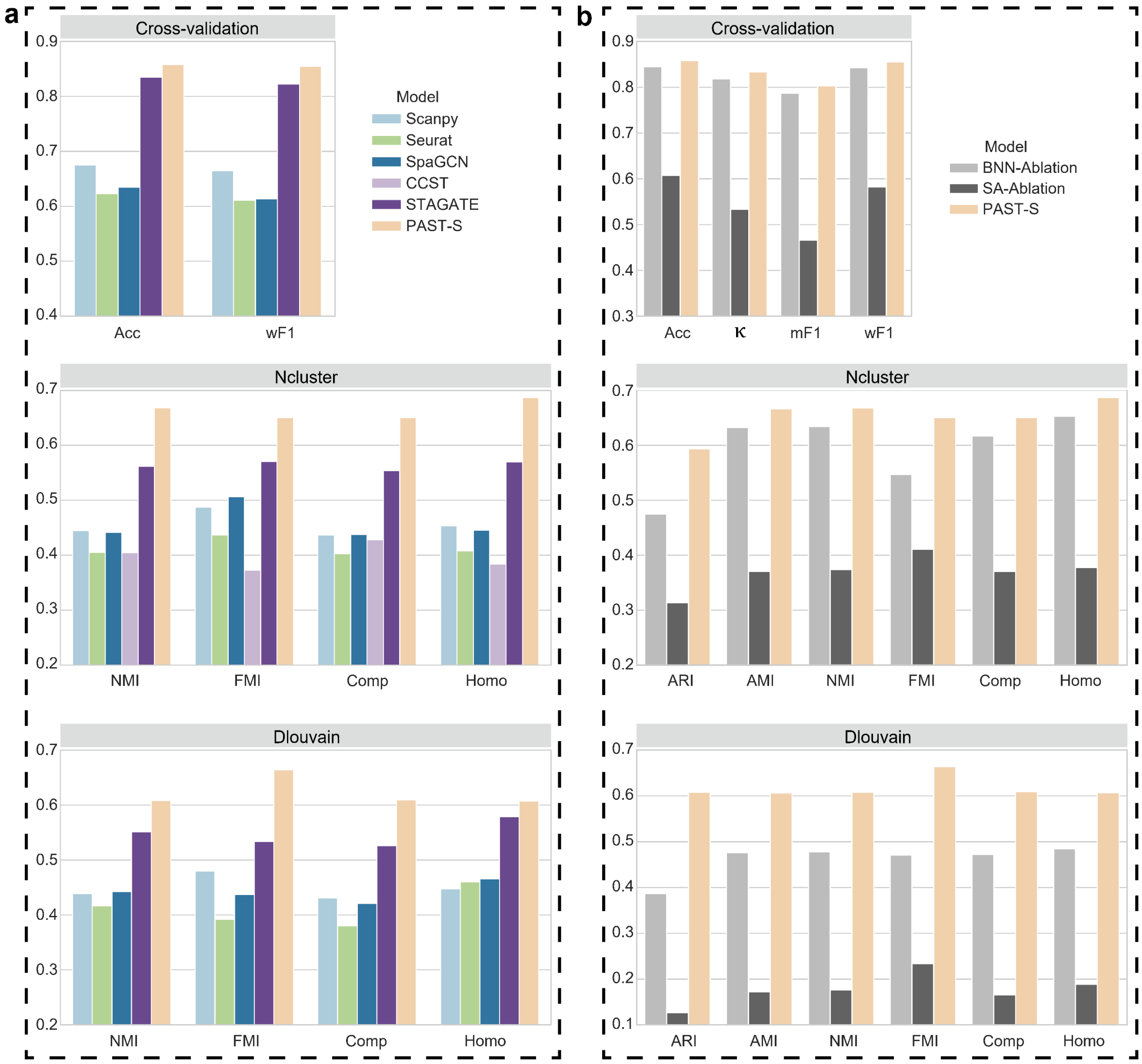

**Fig. S14 | Quantitative performance evaluation of spatial domain characterization on the osmFISH mouse somatosensory cortex (MSC) dataset.** Quantitative evaluation via supervised cross-validation and unsupervised spatial clustering with a specified number of clusters (Ncluster) and with default resolution (Dlouvain). **a**, Evaluation of PAST and other baseline methods. The cross-validation performance was evaluated by the average score of Acc and wF1, respectively, in the 5-fold experiments. The spatial clustering performance was evaluated by NMI, FMI, Comp and Homo. **b**, Evaluation of PAST and ablation models. The cross-validation performance was evaluated by the average score of Acc, κ, mF1 and wF1, respectively, in the 5-fold experiments. The spatial clustering performance was evaluated by ARI, AMI, NMI, FMI, Comp and Homo.

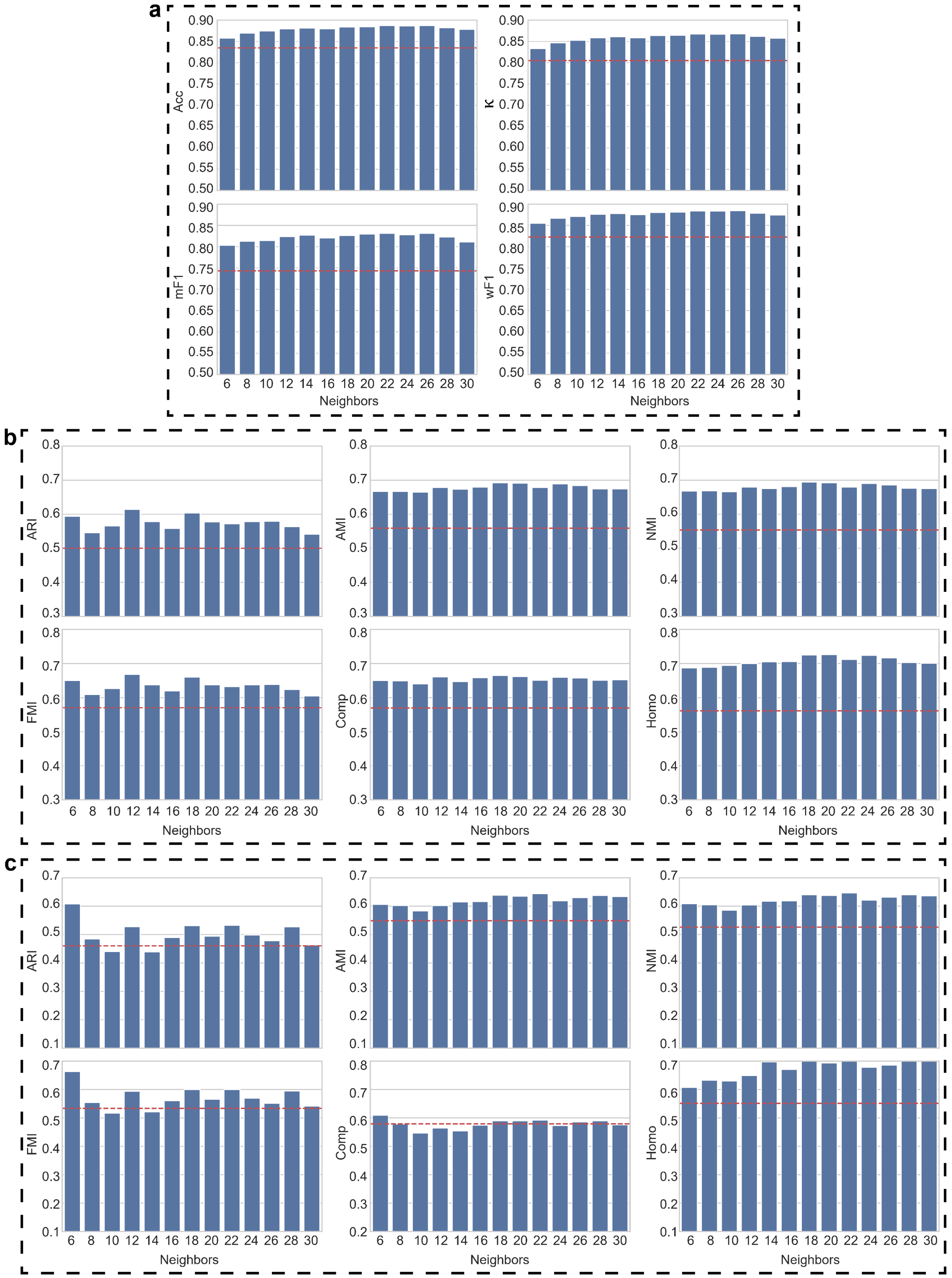

**Fig. S15 | The performance of PAST with different neighbors on osmFISH MSC dataset.** We changed the number of neighbors in self-attention mechanism to explore the robustness of PAST. The red lines denote the second-best method, STAGATE. **a**, Supervised cross-validation evaluated by Acc, κ, mF1 and wF1. Spatial clustering with **b**, a specified number of clusters (Ncluster) or **c**, default resolution (Dlouvain) evaluated by ARI, AMI, NMI, FMI, Comp and Homo.

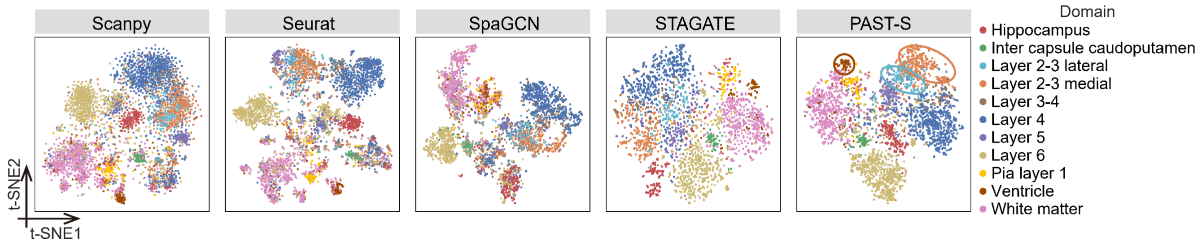

**Fig. S16 | t-SNE Visualization on osmFISH MSC dataset.** The blue, orange and brown circles denote Layer 2-3 lateral, Layer 2-3 medial and Ventricle, respectively.

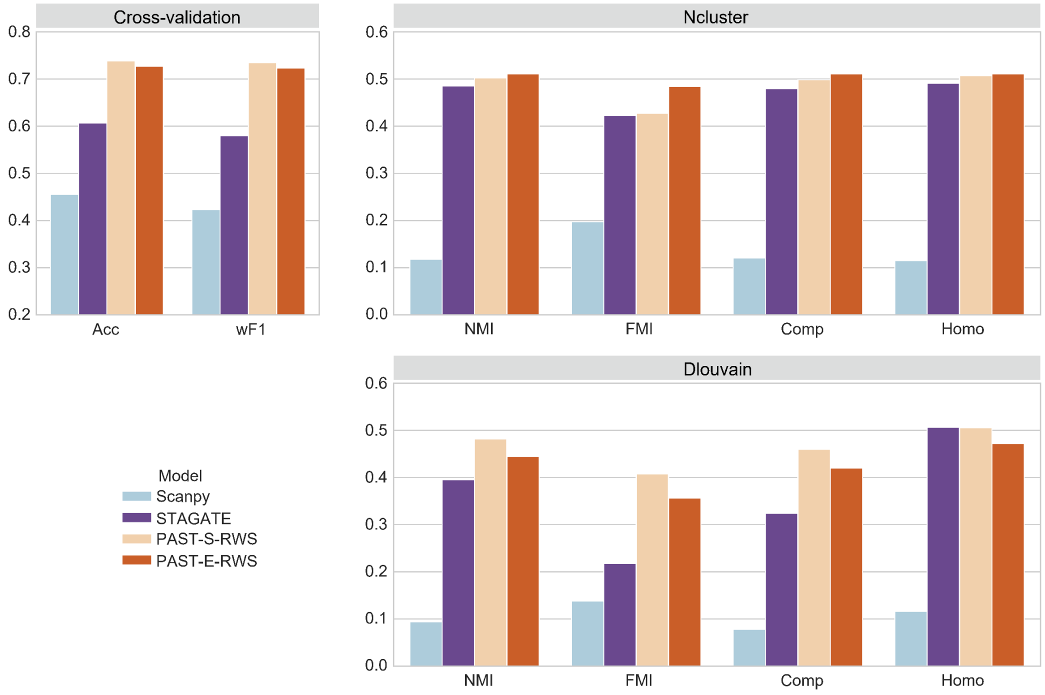

**Fig. S17 | Quantitative performance evaluation of spatial domain characterization on the Stereo-seq mouse olfactory bulb (MOBS1) dataset.** Quantitative evaluation via supervised cross-validation and unsupervised spatial clustering with a specified number of clusters (Ncluster) and with default resolution (Dlouvain). The cross-validation performance was evaluated by the average score of Acc and wF1, respectively, in the 5-fold experiments. The spatial clustering performance was evaluated by NMI, FMI, Comp and Homo.

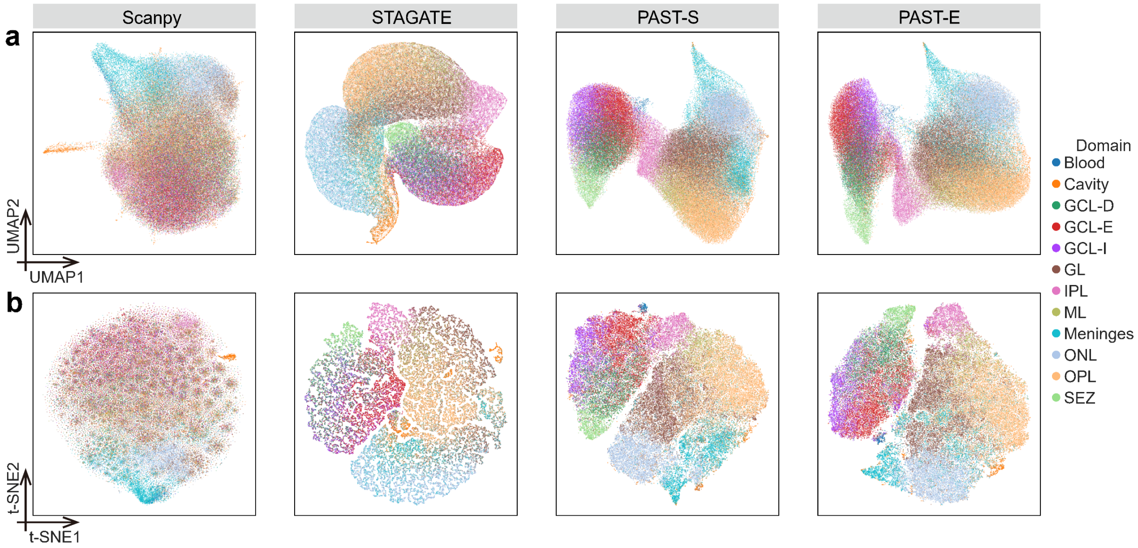

**Fig. S18 | UMAP and t-SNE visualization on Stereo-seq MOBS1 dataset. a**, UMAP visualization and **b**, t-SNE visualization of different methods.

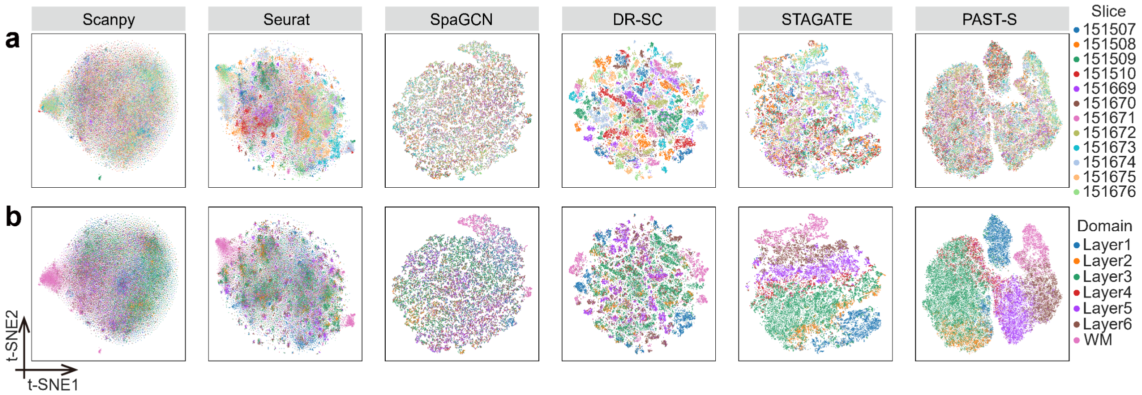

**Fig. S19 | t-SNE visualization of spots in all the DLPFC slices.** We utilized the proposed strategy to obtain the joint embeddings of all spots in DLPFC datasets and then visualized them in t-SNE space colored by **a**, slices and **b**, spatial domains.

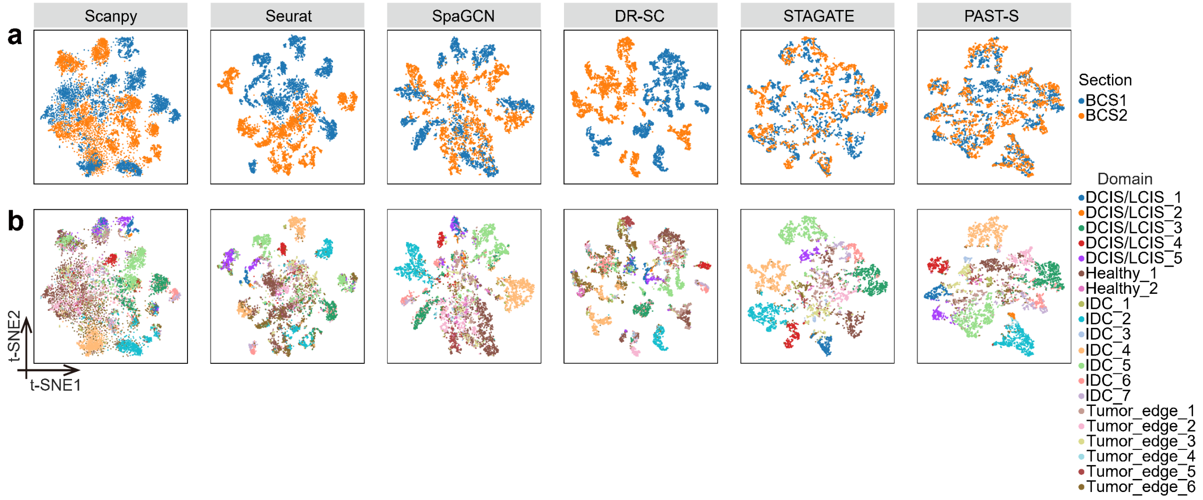

**Fig. S20 | t-SNE visualization of spots in 10x Visium breast cancer sections (BCS1/2).** We utilized the proposed strategy to obtain the joint embeddings of all spots in BCS1/2 datasets and then visualized them in t-SNE space colored by **a**, sections and **b**, spatial domains.

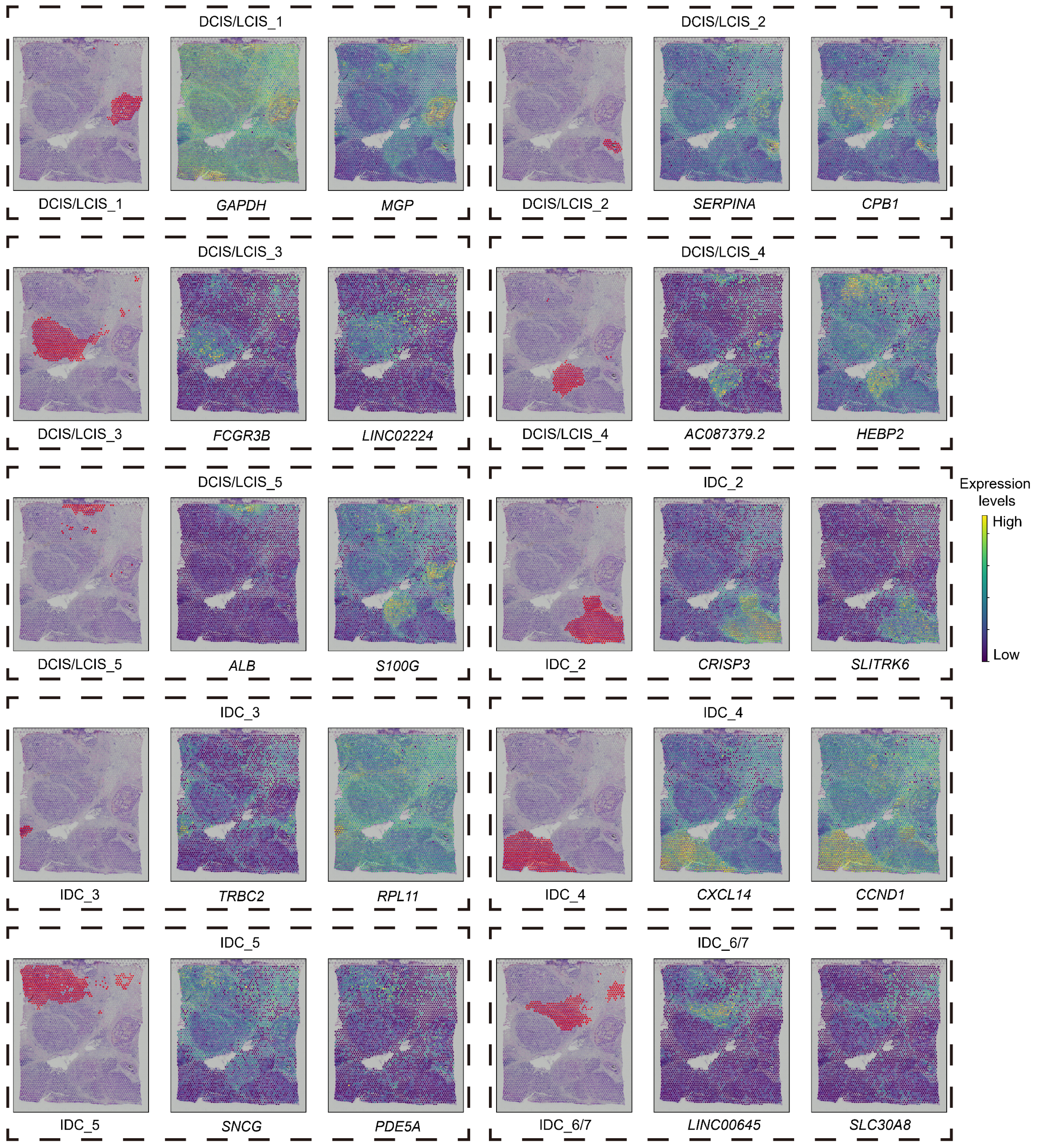

**Fig. S21 | The supervised annotation results of tumor areas based on PAST and the spatial expression patterns of corresponding BCS1-mark genes on BCS2.** We annotated BCS2 based on PAST in a supervised manner and then visualized the spatial expression patterns of BCS1-marker genes on BCS2.

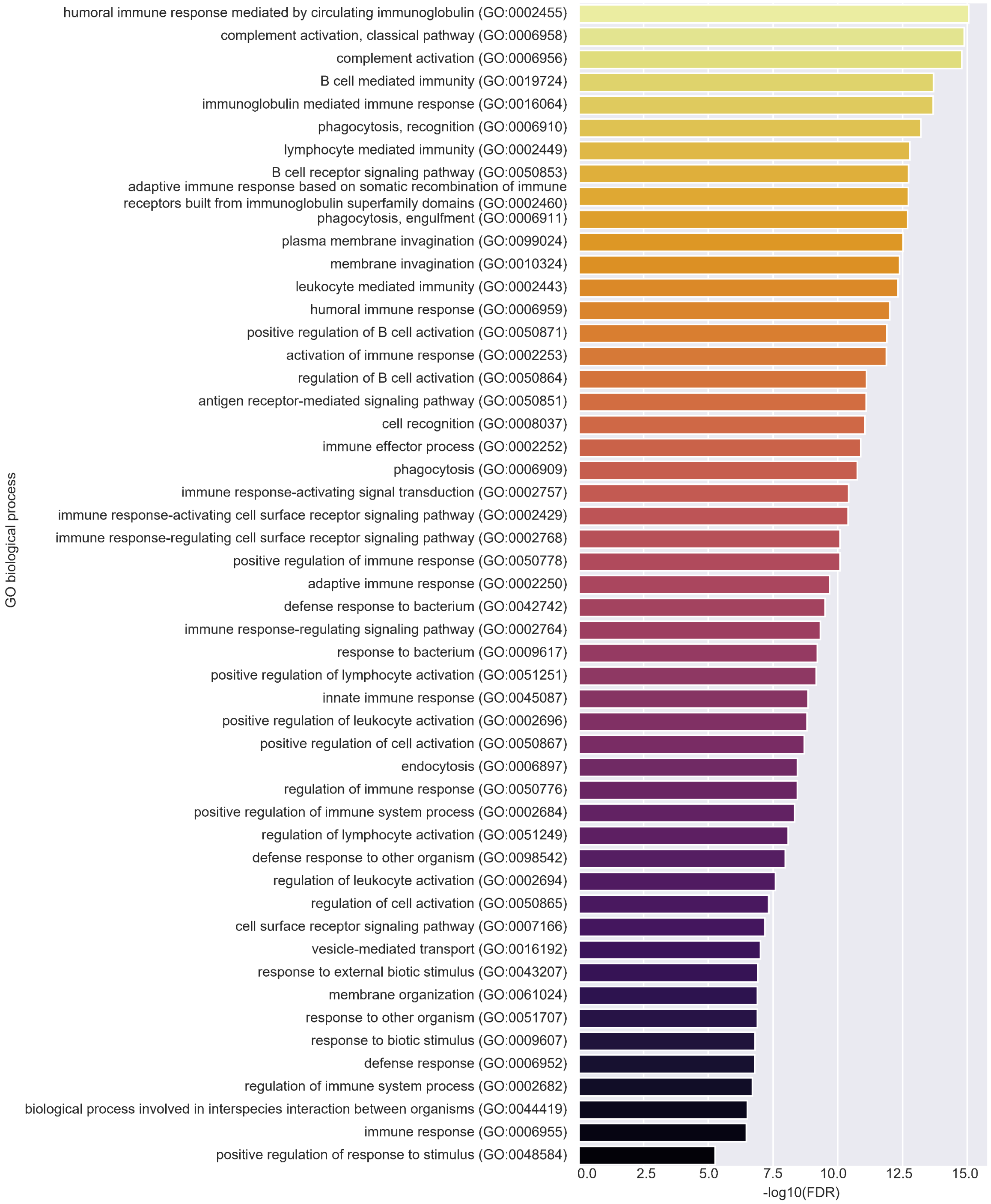

**Fig. S22 | Gene ontology enrichment analysis of Tumor_edge_3 based on the top 10 differential expression genes on BCS2.** We selected the top 10 differential expression genes (DEGs), i.e., *IGHG3, IGKC, IGLC2, IGHG4, IGHG1, IGLC3, C3, IGHA1, IGHM* and *JCHAIN*, with the most significant adjusted *p*-values for the supervised annotated Tumor_edge_3 on BCS2, and then conducted gene ontology (GO) enrichment analysis.

**Table S1. The detailed model structure and hyper-parameters of PAST.** We describe the model structure and corresponding parameter settings in (module name, output dimension) format. Specially, the parameter settings of self-attention module are of (module name, output dimension of [Q, K, V]) format. BNN: Bayesian neural network, FC: fully-connected layer, SA: self-attention module. We utilized Adam optimizer to update the parameters of PAST with initial learning rate of 1e-3 and weight decay of 1e-4. $D$ and $d$ denote the dimension of preprocessed target gene expression matrix and latent space, respectively.

| | |
|---|---|
| **Encoder** | (Target gene expression matrix, $D$) |
| | (BNN,$4d/5$) and (FC, $d/5$ ) |
| | (Concatenation, $d$) |
| | (SA, $[d,d,d]$)×2 |
| **Gaussian distribution** | (FC, $d$) and (FC, $d$) |
| | (Reparameterization trick, $d$) |
| **Decoder** | (SA, $[d,d,d]$)×2 |
| | (FC, $D$) |
| | (Reconstructed target gene expression matrix, $D$) |

**Table S2. Summary of datasets used in this study.** We collected multiple datasets generated by different technologies, including 10x Visium, STARmap, osmFISH and Stereo-seq, and we also collected the corresponding external reference datasets. DLPFC\151xxx denotes all slices except for 151xxx in human DLPFC datasets.

| Target | | | | | | Reference | | | | | |
|---|---|---|---|---|---|---|---|---|---|---|---|
| **Dataset** | **Spots** | **Genes** | **Regions** | **Technology** | **Resolution** | **Dataset** | **Spots** | **Genes** | **Regions** | **Technology** | **Resolution** |
| Human DLPFC 151507 | 4221 | 33538 | 7 | 10x Visium | Non single-cell | DLPFC\151507 | 43108 | 33538 | 7 | 10x Visium | Non single-cell |
| Human DLPFC 151508 | 4381 | 33538 | 7 | 10x Visium | Non single-cell | DLPFC\151508 | 42948 | 33538 | 7 | 10x Visium | Non single-cell |
| Human DLPFC 151509 | 4788 | 33538 | 7 | 10x Visium | Non single-cell | DLPFC\151509 | 42541 | 33538 | 7 | 10x Visium | Non single-cell |
| Human DLPFC 151510 | 4595 | 33538 | 7 | 10x Visium | Non single-cell | DLPFC\151510 | 42734 | 33538 | 7 | 10x Visium | Non single-cell |
| Human DLPFC 151669 | 3636 | 33538 | 5 | 10x Visium | Non single-cell | DLPFC\151669 | 43693 | 33538 | 7 | 10x Visium | Non single-cell |
| Human DLPFC 151670 | 3484 | 33538 | 5 | 10x Visium | Non single-cell | DLPFC\151670 | 43845 | 33538 | 7 | 10x Visium | Non single-cell |
| Human DLPFC 151671 | 4093 | 33538 | 5 | 10x Visium | Non single-cell | DLPFC\151671 | 43236 | 33538 | 7 | 10x Visium | Non single-cell |
| Human DLPFC 151672 | 3888 | 33538 | 5 | 10x Visium | Non single-cell | DLPFC\151672 | 43441 | 33538 | 7 | 10x Visium | Non single-cell |
| Human DLPFC 151673 | 3611 | 33538 | 7 | 10x Visium | Non single-cell | DLPFC\151673 | 43718 | 33538 | 7 | 10x Visium | Non single-cell |
| Human DLPFC 151674 | 3635 | 33538 | 7 | 10x Visium | Non single-cell | DLPFC\151674 | 43694 | 33538 | 7 | 10x Visium | Non single-cell |
| Human DLPFC 151675 | 3566 | 33538 | 7 | 10x Visium | Non single-cell | DLPFC\151675 | 43763 | 33538 | 7 | 10x Visium | Non single-cell |
| Human DLPFC 151676 | 3431 | 33538 | 7 | 10x Visium | Non single-cell | DLPFC\151676 | 43898 | 33538 | 7 | 10x Visium | Non single-cell |
| Mouse primary visual cortex | 1207 | 1020 | 7 | STARmap | Single-cell | Mouse brain coronal | 2688 | 18078 | 15 | 10x Visium | Non single-cell |
| | | | | | | Mouse primary visual cortex | 14249 | 34617 | 23 | scRNA-seq | Single-cell |
| Mouse somatosensory cortex | 4839 | 33 | 11 | osmFISH | Single-cell | - | - | - | - | - | - |
| Mouse olfactory bulb section1 | 107416 | 26145 | 12 | Stereo-seq | Single-cell | Mouse olfactory bulb section2 | 104931 | 23815 | 11 | Stereo-seq | Single-cell |
| Human breast cancer section2 | 3987 | 36601 | NA | 10x Visium | Non single-cell | Human breast cancer section1 | 3798 | 36601 | 20 | 10x Visium | Non single-cell |

**Table S3. Quantitative evaluation of PAST and other methods on the DLPFC datasets.** The median score across all slices of DLPFC datasets in terms of the 16 metrics. Cross validation: 5-fold cross-validation experiments where the latent embeddings were taken as input, the ground-truth labels were taken as output and a support vector machine with default parameters was taken as classifier. Ncluster: spatial clustering with a specified number of clusters. Dlouvain: spatial clustering with default resolution. The numbers in bold represent the best method for given metric in the given scenario.

| Scenario | Metrics | Scanpy | Seurat | SpaGCN | BayesSpace | CCST | DR-SC | STAGATE | PAST-S | PAST-E |
|---|---|---|---|---|---|---|---|---|---|---|
| Cross-validation | Acc | 0.723 | 0.807 | 0.809 | - | - | 0.826 | 0.886 | **0.899** | 0.897 |
| | κ | 0.644 | 0.740 | 0.707 | - | - | 0.778 | 0.850 | 0.866 | **0.867** |
| | mF1 | 0.608 | 0.727 | 0.681 | - | - | 0.762 | 0.818 | **0.870** | **0.870** |
| | wF1 | 0.696 | 0.791 | 0.795 | - | - | 0.819 | 0.875 | **0.898** | 0.896 |
| Ncluster | ARI | 0.198 | 0.290 | 0.352 | 0.403 | 0.452 | 0.454 | 0.515 | 0.529 | **0.551** |
| | AMI | 0.249 | 0.366 | 0.497 | 0.573 | 0.639 | 0.563 | 0.650 | 0.657 | **0.686** |
| | NMI | 0.251 | 0.368 | 0.498 | 0.574 | 0.639 | 0.564 | 0.651 | 0.657 | **0.686** |
| | FMI | 0.371 | 0.443 | 0.503 | 0.535 | 0.593 | 0.573 | 0.600 | 0.617 | **0.665** |
| | Comp | 0.241 | 0.377 | 0.471 | 0.571 | **0.671** | 0.553 | 0.635 | 0.650 | 0.670 |
| | Homo | 0.260 | 0.360 | 0.509 | 0.620 | 0.603 | 0.610 | 0.673 | 0.677 | **0.700** |
| Dlouvain | ARI | 0.182 | 0.259 | 0.240 | - | - | 0.300 | 0.255 | 0.314 | **0.408** |
| | AMI | 0.250 | 0.381 | 0.434 | - | - | 0.499 | 0.515 | 0.546 | **0.602** |
| | NMI | 0.252 | 0.383 | 0.436 | - | - | 0.501 | 0.517 | 0.547 | **0.603** |
| | FMI | 0.371 | 0.383 | 0.353 | - | - | 0.406 | 0.396 | 0.435 | **0.523** |
| | Comp | 0.247 | 0.344 | 0.368 | - | - | 0.425 | 0.422 | 0.466 | **0.533** |
| | Homo | 0.255 | 0.441 | 0.567 | - | - | 0.609 | 0.688 | 0.686 | **0.727** |

**Table S4. Quantitative evaluation of PAST and other methods on the STARmap MPVC dataset.** Cross validation: 5-fold cross-validation experiments where the latent embeddings were taken as input, the ground-truth labels were taken as output and a support vector machine with default parameters was taken as classifier. Ncluster: spatial clustering with a specified number of clusters. Dlouvain: spatial clustering with default resolution. The numbers in bold represent the best method for given metric in the given scenario.

| Scenario | Metrics | Scanpy | Seurat | SpaGCN | CCST | STAGATE | PAST-S | PAST-E-Visium | PAST-E-scRNA |
|---|---|---|---|---|---|---|---|---|---|
| Cross-validation | Acc | 0.665 | 0.665 | 0.654 | - | 0.742 | **0.785** | 0.754 | 0.731 |
| | κ | 0.597 | 0.597 | 0.585 | - | 0.691 | **0.742** | 0.705 | 0.676 |
| | mF1 | 0.625 | 0.622 | 0.556 | - | 0.715 | **0.754** | 0.742 | 0.665 |
| | wF1 | 0.658 | 0.658 | 0.639 | - | 0.738 | **0.778** | 0.751 | 0.720 |
| Ncluster | ARI | 0.296 | 0.288 | 0.348 | 0.503 | 0.388 | **0.564** | 0.553 | 0.533 |
| | AMI | 0.317 | 0.298 | 0.384 | 0.542 | 0.470 | **0.643** | 0.617 | 0.571 |
| | NMI | 0.323 | 0.304 | 0.389 | 0.546 | 0.474 | **0.646** | 0.620 | 0.575 |
| | FMI | 0.413 | 0.403 | 0.451 | 0.591 | 0.496 | **0.644** | 0.634 | 0.610 |
| | Comp | 0.326 | 0.303 | 0.386 | 0.565 | 0.488 | **0.670** | 0.644 | 0.579 |
| | Homo | 0.320 | 0.305 | 0.393 | 0.528 | 0.462 | **0.623** | 0.598 | 0.570 |
| Dlouvain | ARI | 0.303 | 0.291 | 0.342 | - | 0.347 | **0.502** | 0.384 | 0.445 |
| | AMI | 0.317 | 0.301 | 0.383 | - | 0.384 | **0.607** | 0.493 | 0.507 |
| | NMI | 0.322 | 0.307 | 0.389 | - | 0.389 | **0.610** | 0.498 | 0.512 |
| | FMI | 0.415 | 0.405 | 0.444 | - | 0.449 | **0.580** | 0.484 | 0.531 |
| | Comp | 0.321 | 0.306 | 0.377 | - | 0.385 | **0.597** | 0.493 | 0.497 |
| | Homo | 0.323 | 0.308 | 0.401 | - | 0.394 | **0.624** | 0.503 | 0.528 |

**Table S5. Quantitative evaluation of PAST and other methods on the osmFISH MSC dataset.** Cross validation: 5-fold cross-validation experiments where the latent embeddings were taken as input, the ground-truth labels were taken as output and a support vector machine with default parameters was taken as classifier. Ncluster: spatial clustering with a specified number of clusters. Dlouvain: spatial clustering with default resolution. The numbers in bold represent the best method for given metric in the given scenario.

| Scenario | Metrics | Scanpy | Seurat | SpaGCN | CCST | STAGATE | PAST-S |
|---|---|---|---|---|---|---|---|
| Cross-validation | Acc | 0.675 | 0.623 | 0.635 | - | 0.835 | **0.858** |
| | κ | 0.621 | 0.564 | 0.574 | - | 0.805 | **0.834** |
| | mF1 | 0.569 | 0.510 | 0.475 | - | 0.743 | **0.803** |
| | wF1 | 0.665 | 0.611 | 0.614 | - | 0.823 | **0.855** |
| Ncluster | ARI | 0.405 | 0.346 | 0.426 | 0.245 | 0.500 | **0.594** |
| | AMI | 0.442 | 0.402 | 0.439 | 0.402 | 0.559 | **0.667** |
| | NMI | 0.445 | 0.405 | 0.442 | 0.405 | 0.561 | **0.668** |
| | FMI | 0.487 | 0.437 | 0.506 | 0.373 | 0.570 | **0.650** |
| | Comp | 0.437 | 0.403 | 0.438 | 0.428 | 0.553 | **0.650** |
| | Homo | 0.453 | 0.408 | 0.445 | 0.384 | 0.570 | **0.687** |
| Dlouvain | ARI | 0.397 | 0.303 | 0.352 | - | 0.460 | **0.608** |
| | AMI | 0.436 | 0.412 | 0.439 | - | 0.549 | **0.607** |
| | NMI | 0.439 | 0.417 | 0.442 | - | 0.551 | **0.608** |
| | FMI | 0.480 | 0.392 | 0.437 | - | 0.534 | **0.664** |
| | Comp | 0.431 | 0.380 | 0.421 | - | 0.526 | **0.609** |
| | Homo | 0.447 | 0.460 | 0.466 | - | 0.579 | **0.607** |

**Table S6. Quantitative evaluation of PAST and other methods on the Stereo-seq MOBS1 dataset.** Cross validation: 5-fold cross-validation experiments where the latent embeddings were taken as input, the ground-truth labels were taken as output and a support vector machine with default parameters was taken as classifier. Ncluster: spatial clustering with a specified number of clusters. Dlouvain: spatial clustering with default resolution. The numbers in bold represent the best method for given metric in the given scenario.

| Scenario | Metrics | Scanpy | STAGATE | PAST-S | PAST-E |
|---|---|---|---|---|---|
| Cross-validation | Acc | 0.456 | 0.607 | **0.739** | 0.727 |
| | κ | 0.363 | 0.548 | **0.701** | 0.688 |
| | mF1 | 0.382 | 0.513 | **0.704** | 0.700 |
| | wF1 | 0.423 | 0.580 | **0.735** | 0.723 |
| Ncluster | ARI | 0.073 | 0.345 | 0.350 | **0.413** |
| | AMI | 0.118 | 0.485 | 0.503 | **0.511** |
| | NMI | 0.118 | 0.485 | 0.503 | **0.511** |
| | FMI | 0.197 | 0.423 | 0.428 | **0.484** |
| | Comp | 0.121 | 0.480 | 0.499 | **0.511** |
| | Homo | 0.115 | 0.491 | 0.507 | **0.511** |
| Dlouvain | ARI | 0.061 | 0.132 | **0.334** | 0.278 |
| | AMI | 0.093 | 0.395 | **0.481** | 0.444 |
| | NMI | 0.094 | 0.395 | **0.482** | 0.444 |
| | FMI | 0.138 | 0.217 | **0.407** | 0.356 |
| | Comp | 0.078 | 0.324 | **0.460** | 0.420 |
| | Homo | 0.116 | **0.507** | 0.506 | 0.472 |
